# Structural insights into the assembly of gp130 family cytokine signaling complexes

**DOI:** 10.1101/2022.06.30.496838

**Authors:** Yi Zhou, Panayiotis E. Stevis, Jing Cao, Kei Saotome, Jiaxi Wu, Arielle Glatman Zaretsky, Sokol Haxhinasto, George D. Yancopoulos, Andrew J. Murphy, Mark W. Sleeman, William C. Olson, Matthew C. Franklin

## Abstract

The gp130 family cytokine signaling complexes have limited structural information despite their crucial roles in various cellular processes. We determined cryo-EM structures of several complexes of this family, containing full ectodomains of both signaling receptors bound to their respective ligands CNTF, CLCF1, LIF, IL-27, and IL-6. Our structures reveal that gp130 serves as a central receptor by engaging Site 2 of CNTF, CLCF1, LIF, and IL-6, and Site 3 of IL-27 and IL-6. The acute bends at both signaling receptors in all complexes bring the membrane-proximal domains to a ~30 Å range but with distinct distances and orientations, which might determine biological specificities of these cytokines. We also reveal how CLCF1 engages its secretion chaperone CRLF1. Our data provide valuable insights for therapeutically targeting gp130-mediated signaling.

## Introduction

Glycoprotein 130 (gp130) is a signaling receptor for Interleukin 6 (IL-6) family cytokines (or gp130 family cytokines), including IL-6, Ciliary Neurotrophic Factor (CNTF), Cardiotrophin Like Cytokine Factor 1 (CLCF1), Leukemia Inhibitory Factor (LIF), Oncostatin M (OSM), Cardiotrophin-1 (CT-1), IL-11, IL-27, IL-35, and IL-39 (*1*). These cytokines share a canonical four-helix bundle structure in which four major helices (termed helices A-D) are linked by three loops (termed AB, BC, and CD loops), as well as three conserved receptor binding epitopes (termed Sites 1-3). Signals induced by these cytokines play critical and diverse roles in regulation of various cellular processes, including inflammatory and immune responses, embryonic development, neuronal and liver regeneration, and hematopoiesis, while dysregulation of these signals leads to a variety of diseases and cancers (*2*). IL-6 and IL-11 signal through gp130 homodimerization, while other cytokines require another ‘tall’ signaling receptor such as LIF receptor (LIFR) or IL-27 receptor subunit alpha (IL-27Rα) which forms a heterodimer with gp130 for signal transduction (*1*). The extracellular domains (ECDs) of gp130 include an N-terminal Ig-like domain (D1), a cytokine-binding homology region (CHR, D2D3), and three membrane-proximal fibronectin type III domains (FNIII, D4–D6). LIFR and IL-27Rα ECDs share a similar domain arrangement at the C-terminus as gp130, but LIFR has an additional CHR preceding the Ig-like domain while IL-27Rα does not have the N-terminal Ig-like domain.

IL-6 forms a symmetric 2:2:2 signaling complex with gp130 and IL-6 receptor subunit alpha (IL-6Rα) (*3*). CNTF binds to CNTF receptor subunit alpha (CNTFRα) first and then assembles into a 1:1:1:1 quaternary signaling complex with gp130 and LIFR (*4*, *5*). CLCF1 was also found to interact with CNTFRα and signal via the gp130-LIFR heterodimer analogous to CNTF, and Cytokine Receptor Like Factor 1 (CRLF1) was shown to chaperone the secretion of CLCF1 (*6*). LIF forms a 1:1:1 tripartite complex with gp130 and LIFR and the signal transduction does not require a non-signaling alpha receptor (*7*). IL-27 is a heterodimeric cytokine of p28 and Epstein-Barr Virus Induced gene 3 (EBI3) and signals though gp130 and IL-27Rα by forming a 1:1:1:1 quaternary complex (*8*).

It has been proposed that the membrane-proximal domains of the two signaling receptors are brought into close proximity upon assembly of the gp130 family cytokine signaling complexes, which allows trans-phosphorylation of Janus kinases (JAKs) bound to the intracellular domains (ICDs) of the two receptors (*9*, *10*). Specific cytoplasmic tyrosine-containing motifs in these receptors are phosphorylated by JAKs and consequently serve as docking sites for recruitment and activation of the signal transducers and activators of transcription (STATs), which eventually translocate to the nucleus to regulate gene expression (*11*).

Despite the importance of the gp130 family cytokine signaling complexes in diverse cellular processes, the structures of these complexes are not well characterized due to their high flexibility and instability. The IL-6 signaling complex is the best characterized with a 3.65 Å crystal structure of the assembly core region and a low-resolution negative stain electron microscopy (EM) map of the complex with full gp130 ectodomains (*3*, *10*). A low-resolution negative stain EM map was also reported for the CNTF signaling complex, but the assembly details are not clear (*9*). Two other studies have revealed how human gp130 D2D3 and mouse LIFR D1-D5 engage LIF (*7*, *12*). Additionally, a cryo-EM structure of human IL-27 signaling complex assembly core region was reported most recently (*13*). However, the overall architectures of the LIF and IL-27 signaling complexes remain unknown. Moreover, there is no structural information for the CLCF1 signaling complexes and how CLCF1 engages its chaperone CRLF1 is elusive.

We have determined cryogenic electron microscopy (cryo-EM) structures of the signaling complexes for CNTF, CLCF1, LIF, and IL-27 at sub-4 Å resolution using full ectodomains of both signaling receptors in these complexes. We have also obtained a 3.22 Å cryo-EM structure for the IL-6 signaling complex using detergent-solubilized gp130 containing transmembrane domain and intracellular Box1/Box2 motifs where JAKs bind. Our structures reveal that gp130 serves as a central receptor by engaging Site 2 of CNTF, CLCF1, LIF, and IL-6, and Site 3 of IL-27 and IL-6. The acute bends at both signaling receptors in these complexes bring the juxtamembrane domains to a ~30 Å range but with distinct distances and orientations, which might determine biological specificities of the cytokines. Additionally, we have solved a 3.40 Å cryo-EM structure of the CRLF1-CLCF1-CNTFRα complex, which exhibits an unexpected 2:2:2 stoichiometry. CLCF1 Site 2 and Site 3 are engaged by two CRLF1 molecules analogous to how they are engaged by gp130 and LIFR in the CLCF1 signaling complex. Our results have provided valuable insights into the assembly and signaling mechanisms of gp130 family cytokine-receptor complexes.

## Results

### Structural characterization of the CNTF, CLCF1, and LIF signaling complexes

Using cryo-EM, we obtained structures of the human CNTF, CLCF1 and LIF signaling complexes, which exhibit similar structural architectures and assembly mechanisms (Figs. 1 and 2). In these complexes, Site 2 and Site 3 of the ligands bind to gp130 and LIFR, respectively, while Site 1 is occupied by CNTFRα or left empty.

**Fig. 1.**
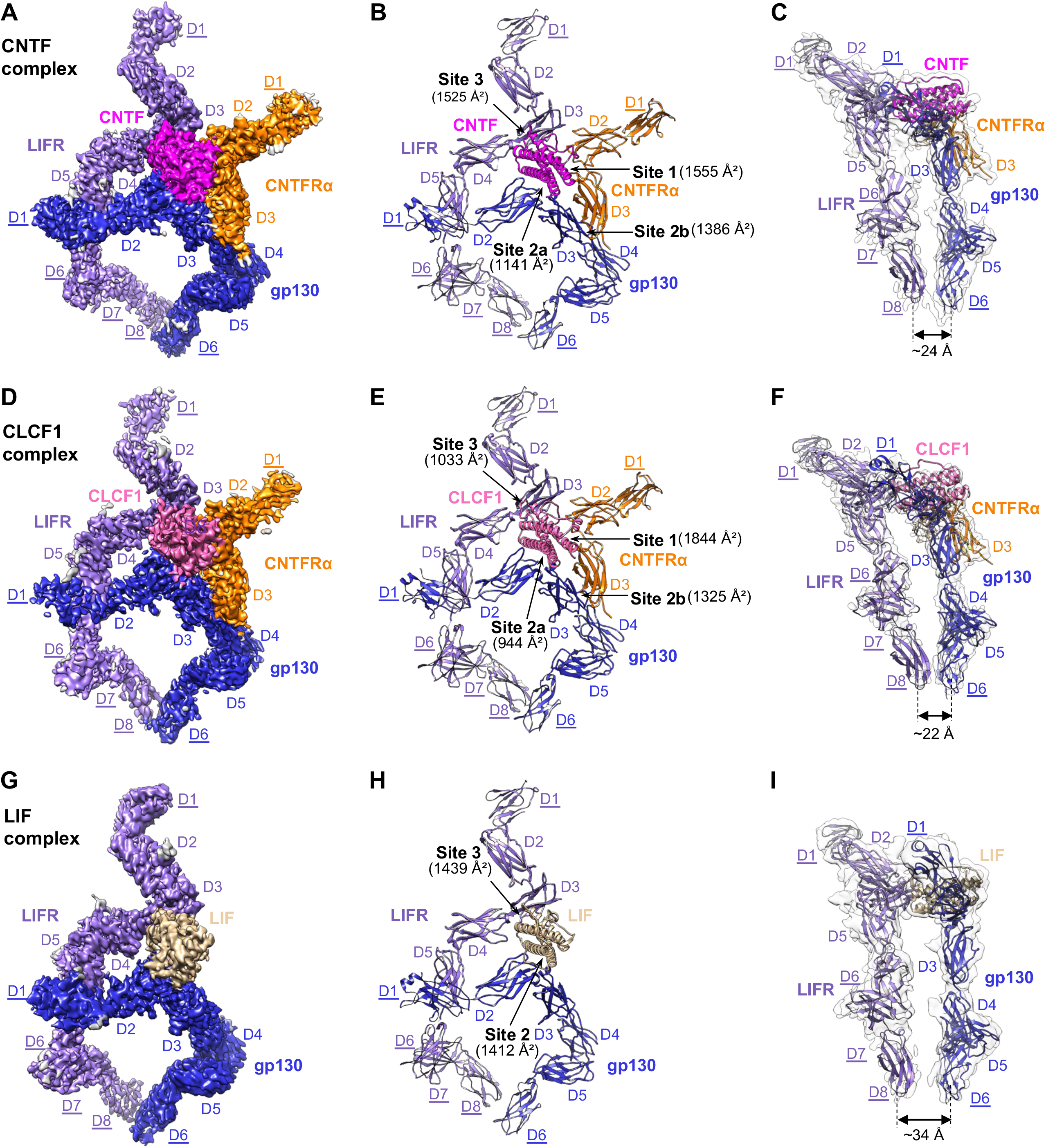
Cryo-EM structures of the CNTF complex, CLCF1 complex and LIF complex. (A-C) The CNTF signaling complex is shown in three representations: colored cryo-EM density map (A), ribbon representation of the model (B), and side view of the model in the transparent EM density map, low-pass filtered to better show density in the peripheral domains (C). (D-F) The CLCF1 signaling complex is shown similarly to panels A-C. (G-I) The LIF signaling complex is shown similarly to panels A-C. Receptor domains that were only rigid-body refined against the density are underlined. The interaction interfaces with corresponding buried surface areas calculated by PDBePISA (*37*) are indicated by arrows in panels B, E, and H. Approximate distances between the bottom centers of LIFR and gp130 juxtamembrane domains in each complex are shown in panels C, F, and I.

**Fig. 2.**
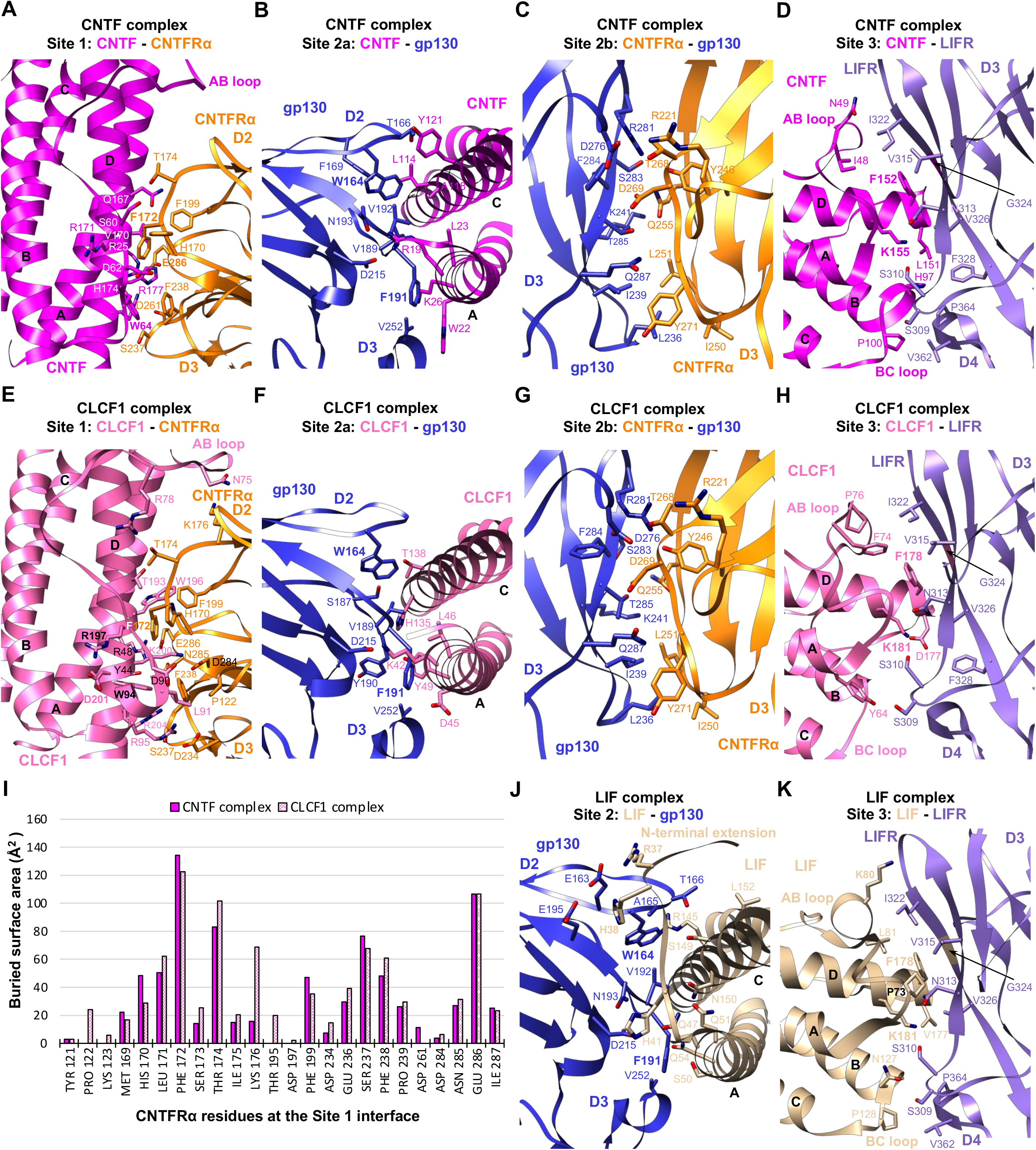
Interaction interfaces of the CNTF complex, CLCF1 complex and LIF complex. (A-D) The binding interfaces of the CNTF complex at Site 1 (A), Site 2a (B), Site 2b (C) and Site 3 (D) as indicated in Fig. 1B are shown in cartoon form, with residues involved in binding shown in stick representation. (E-H) The CLCF1 complex binding interfaces at Site 1 (E), Site 2a (F), Site 2b (G) and Site 3 (H) as indicated in Fig. 1E are shown in the same representation as panels A-D. (I) Histogram of buried surface area for CNTFRα residues at the Site 1 interface in the CNTF and CLCF1 signaling complexes. (J-K) Details of the LIF complex binding interfaces at Site 2 (J) and Site 3 (K) as indicated in Fig. 1H.

### Cryo-EM structure of the CNTF signaling complex

We reconstituted the quaternary CNTF signaling complex using full ectodomains of gp130 and LIFR, and a fusion protein of CNTFRα and CNTF linked by a flexible linker. The complex was characterized by single-particle cryo-EM, generating a density map with a global resolution of 3.03 Å (fig. S1, A to F). This map has well resolved density around the interaction core region, including CNTF, CNTFRα D2D3, gp130 D2-D5, and LIFR D2-D5, permitting model building and full refinement of this region. Due to the flexible nature of the receptors, the local resolution at the distal ends of the receptors is lower. The membrane-proximal FNIII domains (D6-D8) of LIFR have fragmented density, suggesting high heterogeneity induced by flexibility of this region. A subset of particles was further identified by heterogeneous refinement, yielding another map with lower global resolution (3.37 Å), but improved density for LIFR D6-D8 (Fig. 1A; fig. S1, G and H). The new map was used for placement and rigid-body refinement of models for the receptor distal domains, which were derived from published structures (gp130 D1 (*3*), gp130 D6 (*2*), and LIFR D1 (*9*)), or predicted by AlphaFold (LIFR D6-D8) (*14*). Additionally, to help building the model for CNTFRα D1, we determined a 2.93Å cryo-EM structure of CNTFRα full ectodomain with the help of two antibody Fab fragments bound to CNTFRα D1 and D2, respectively (fig. S2). The CNTFRα D1 model derived from this structure was also rigid-body refined against the 3.37 Å CNTF signaling complex map. Combining all of this, we generated a complete model of the CNTF signaling complex with full ECDs (Fig. 1B). Intriguingly, the acute bends of gp130 at D4D5 and LIFR at D6D7 bring the bottom centers of the receptor juxtamembrane domains to ~24 Å apart (Fig. 1C).

CNTF Site 1 is occupied by CNTFRα, similar to how IL-6 is engaged by IL-6Rα (*3*). CNTFRα D2D3 adopts an elbow-like conformation and holds CNTF in its hinge region by interacting with CNTF helices A and D, and AB loop (Fig. 1B and Fig. 2A). The interface is enriched with charged residues, including CNTFRα^D261, E286^ and CNTF ^R25, H174, R177^, which mediate a network of hydrogen bonds and salt bridge interactions. The interface is centered around CNTFRα^F172^, which is surrounded by W64, V170, R171, and H174 of CNTF. The C-terminus of CNTF AB loop is held in position through hydrophobic interactions between CNTF^W64^ and CNTFRα^F172, F238^. Notably, F172 and E286 in CNTFRα are two residues that contribute the two highest buried surface areas (134 Å^2^ and 107 Å^2^, respectively), consistent with their key roles in ligand binding (*15*). The importance of CNTF^W64^ in the interaction is also supported by a previous mutagenesis study (*16*).

On the other side of Site 1, CNTF helices A and C are captured by the elbow region of gp130 CHR (D2D3), forming the Site 2a interface (Figs. 1B and 2B). Similar to the way gp130 engages LIF (*12*) and IL-6 (*3*), gp130^F191^ contributes the largest fraction of buried surface area (119 Å^2^) at the gp130-CNTF interface by inserting into a hydrophobic pocket formed by W22, L23, and the hydrophobic portions of R19 and K26 on helix A of CNTF. CNTF^R19^ is also coordinated by gp130D^215^ to form hydrogen bond and salt bridge at the center of the interface. CNTF helix C is held in position by gp130^W164^ packing against the middle of the helix. While the interactions are predominantly mediated by gp130 D2, V252 from D3 interacts with W22 and K26 at the middle of CNTF helix A, which likely improves the binding.

CNTFRα D3 leans against gp130 D3 to make a “stem-stem” Site 2b interaction, contributing an additional 1,386 Å^2^ buried surface area to the composite Site 2 (Fig. 1B), which likely increases the overall binding affinity. Consistent with this, CNTF is not able to initiate signaling in the absence of CNTFRα (*5*). The Site 2b interface is dominated by hydrophilic interactions and centered around CNTFRα^D269^ which pairs with gp130^R281, T285^ (Fig. 2C). A hydrophobic patch of CNTFRα residues (I250, L251, Y271) further anchors the distal end of its D3 to gp130 by engaging L236 and I239 of gp130.

LIFR contacts the posterior end of the CNTF four-helix bundle, including the N-terminus of AB loop and helix D, C-terminus of helix B, and the short BC loop, to form the Site 3 interface where 27 LIFR residues and 25 CNTF residues together bury a total of 1,525 Å^2^ surface area (Figs. 1B and 2D). The interactions are predominately mediated by LIFR D3 (Ig) domain, with the N-terminal loop of D4 serving as a supporting binding site. Two residues at the N-terminus of CNTF helix D, F152 and K155, play crucial roles in the binding, with additional interactions mediated by the packing of hydrophobic patches at the CNTF AB and BC loops against LIFR D3 and D4, respectively. The aromatic ring of CNTF^F152^ makes a π-stacking against the peptide bond of LIFR^G324^, which is sitting at the bottom of a hydrophobic cavity formed by N313, V315, I322, and V326 of LIFR. CNTF^K155^ is coordinated by LIFR ^S310, N313^ to form hydrogen bonds. Supporting these observations, CNTF F152 and K155 have been shown to be essential for binding to LIFR in a mutagenesis study (*17*). Notably, CNTF F152 and K155 form a F*XX*K motif that is evolutionarily conserved among other IL-6 family cytokines that bind to LIFR. The equivalent residues in LIF and CLCF1 are F178 and K181. It was reported that F178 and K181 in human LIF (hLIF) engage mouse LIFR (mLIFR) in a highly similar manner (*7*).

### Cryo-EM structure of the CLCF1 signaling complex

CLCF1 signals through the same receptors as CNTF. Using full ectodomains of gp130 and LIFR, and a fusion protein of CNTFRα and CLCF1 linked by a flexible linker, we obtained a 3.90 Å cryo-EM map of the CLCF1 quaternary signaling complex, which has ~3.40 Å local resolution around the interaction core region permitting model building (Fig. 1, D and E; fig. S3, A to E). Density at the receptor distal ends was sufficient for domain placement and rigid-body refinement as with the CNTF complex. The bottom centers of LIFR and gp130 juxtamembrane domains are positioned ~22 Å apart (Fig. 1F), comparable to that seen in the CNTF complex.

Similar to CNTF, CLCF1 Site 1 is captured by the elbow region of CNTFRα (Fig. 1E). CNTFRα^F172^ also serves as the anchor point of the CNTFRα-CLCF1 interface by engaging T193, W196, R197, and K200 of CLCF1 (Fig. 2E). Multiple electrostatic interactions are observed between positively charged CLCF1 residues (R48, R95, K200) and negatively charged CNTFRα residues (D234, D284, E286). Notably, each of the key CNTFRα residues, including F172, T174, and E286, contributes significant binding interactions to both CNTF and CLCF1 (Fig. 2I). It has been reported that W94, R197 and D201 of CLCF1 are critical for binding to CNTFRα (*18*). A CLCF1^R197L^ mutation allele was also identified in patients and the mutated protein failed to bind to CNTFRα (*19*). These observations align well with our structure which suggests that W94 plays an important role in packing the C-terminus of the CLCF1 AB loop against CNTFRα though a hydrophobic interaction with CNTFRα F238 (Fig. 2E). The hydrophobic portion of CLCF1^R197^ also directly engages CNTFRα by leaning against CNTFRα^F172^. Interestingly, CLCF1D^201^ does not directly contact CNTFRα; however, it forms a hydrogen bond with CLCF1^W94^ and likely holds W94 in position to interact with CNTFRα.

Like CNTF, CLCF1 binds to gp130 D2D3 to form the Site 2a interface (Fig. 1E). Key gp130 residues engaging CNTF maintain their roles in the CLCF1 complex, with F191 engaging K42, D45, L46, and Y49 of CLCF1 Helix A, D215 forming a hydrogen bond with CLCF1^K42^ at the center of the interface, and W164 capturing the middle portion of CLCF1 Helix C (Fig. 2F).

The CLCF1 complex shares the same Site 2b interface as the CNTF complex where CNTFRα D3 is docked into a wide cavity of gp130 D3 to stabilize the binding of CLCF1 to gp130 (Figs. 1E and 2G). Similarly, Site 2b (1,325 Å^2^) has a larger buried surface area than Site 2a (944 Å^2^) in the CLCF1 complex and significantly increases interactions at the composite Site 2, which may explain why CNTFRα is indispensable for CLCF1 signaling (*6*).

CLCF1 also binds to LIFR D3 to form the Site 3 interface. Notably, 72% of CLCF1 residues contributing to the packing of this interface are non-polar, which is much higher than the percentage of non-polar CNTF residues at the CNTF-LIFR interface (44%). Despite this difference in residue polarity, LIFR captures CLCF1 in a similar manner to how it engages CNTF. Like F152 and K155 of CNTF, the two conserved CLCF1 residues F178 and K181 also serve as key anchor points engaging LIFR residues (S310, N313, V315, I322, G324, and V326) (Fig. 2H), consistent with previous mutagenesis studies (*18*, *20*). CLCF1 helix B does not make significant contacts with the N-terminal loop of LIFR D4 as CNTF helix B does, because CLCF1 helix B is slightly shorter than CNTF helix B at the C-terminus. In addition, fewer hydrogen bonds are formed at the CLCF1-LIFR interface compared with the CNTF-LIFR interface due to lower percentage of polar/charged residues at CLCF1 Site 3. These two factors together may have led to a significantly lower buried surface area of the CLCF1-LIFR interface (1,033 Å^2^) than the CNTF-LIFR interface (1,525 Å^2^).

### Cryo-EM structure of the LIF signaling complex

LIF signals via gp130 and LIFR in the absence of a non-signaling alpha receptor. The overall architecture of this tripartite signaling complex remains unknown. We therefore solved a 3.54 Å cryo-EM map of the complex containing full ectodomains of gp130 and LIFR. As with the CNTF and CLCF1 complexes, this map’s higher resolution at the interaction core region permitted full model building and refinement, while the density at the distal ends of the receptors was sufficient for domain placement and rigid-body refinement (Fig. 1, G and H; fig. S3, F to K).

The gp130-LIF interface has a larger buried surface area (1,412 Å^2^) than the gp130-CNTF and gp130-CLCF1 interfaces (Fig. 1), which is mostly caused by gp130 engaging LIF N-terminal extension (residues 34-42) preceding helix A (Fig. 2J). Consistent with the crystal structure of gp130 D2D3 in complex with LIF (*12*), the LIF N-terminal extension serves as a “doorstop” to hold gp130 in position and creates additional interaction interface, which might be the reason why LIF does not need a non-signaling alpha receptor. Of the 21 LIF residues forming the contacting surface, R37 and H38 from the N-terminal extension contribute the two highest fractions of buried surface area through electrostatic interactions with gp130^E163, E195^. In contrast, similar N-terminal extension with a rigid conformation is not seen in CNTF or CLCF1. W164 and F191 of gp130 also engage LIF analogous to how they interact with CNTF and CLCF1. Interestingly, while gp130 largely covers the middle region of helices A and C of CNTF and CLCF1, it is positioned towards the N-terminus of LIF helix A, leading to an increased distance between the bottom centers of gp130 D6 and LIFR D8 in the LIF complex (~34 Å) (Fig. 1I).

Like CNTF and CLCF1, LIF also engages LIFR via the conserved F*XX*K motif at Site 3 (Fig. 2K). The way hLIFR residues (S310, N313, V315, I322, G324, and V326) interact with hLIF^F178, K181^ is also observed in the hLIF/mLIFR D1-D5 complex crystal structure (*7*), consistent with the cross-reactivity of hLIF with both hLIFR and mLIFR. LIF is primarily docked onto the saddle-shaped LIFR D3 through its helix D and AB loop, with additional contacts made between LIF BC loop and the N-terminal loop of LIFR D4. In addition to the key residues F178 and K181 in LIF helix D, P73 and K80 in the AB loop also make extensive contacts with LIFR. Consistent with our structure, a previous alanine substitution study has proven the importance of these LIF residues in LIFR binding (*21*).

### Structural characterization of the CRLF1-CLCF1-CNTFRα complex

The secretion of CLCF1 was shown to require its chaperone CRLF1 (*6*). However, how CRLF1 engages CLCF1 is unclear. It was reported that CRLF1 forms a tripartite complex with CLCF1 and CNTFRα and promotes CLCF1 binding to CNTFRα (*22*). We obtained a 3.40 Å cryo-EM structure of the human CRLF1-CLCF1-CNTFRα complex (Fig. 3; fig. S4, A to F). Surprisingly, the complex is a 2:2:2 hexamer with 2-fold symmetry (Fig. 3, A and B). The two CNTFRα molecules bind to Site 1 of the two CLCF1 ligands without contacting CRLF1, indicating that CNTFRα may not be essential for CLCF1-mediated CRLF1 dimerization. Consistent with this, we solved a 3.45 Å cryo-EM structure of the CRLF1-CLCF1 complex, showing that CLCF1 and CRLF1 are able to form a symmetric 2:2 tetramer in the absence of CNTFRα (fig. S4, G to K). Interestingly, the two CRLF1 molecules captures two CLCF1 ligands analogous to how two gp130 receptors engages two viral IL-6 (vIL-6) ligands (fig. S4, K and L) (*23*).

**Fig. 3.**
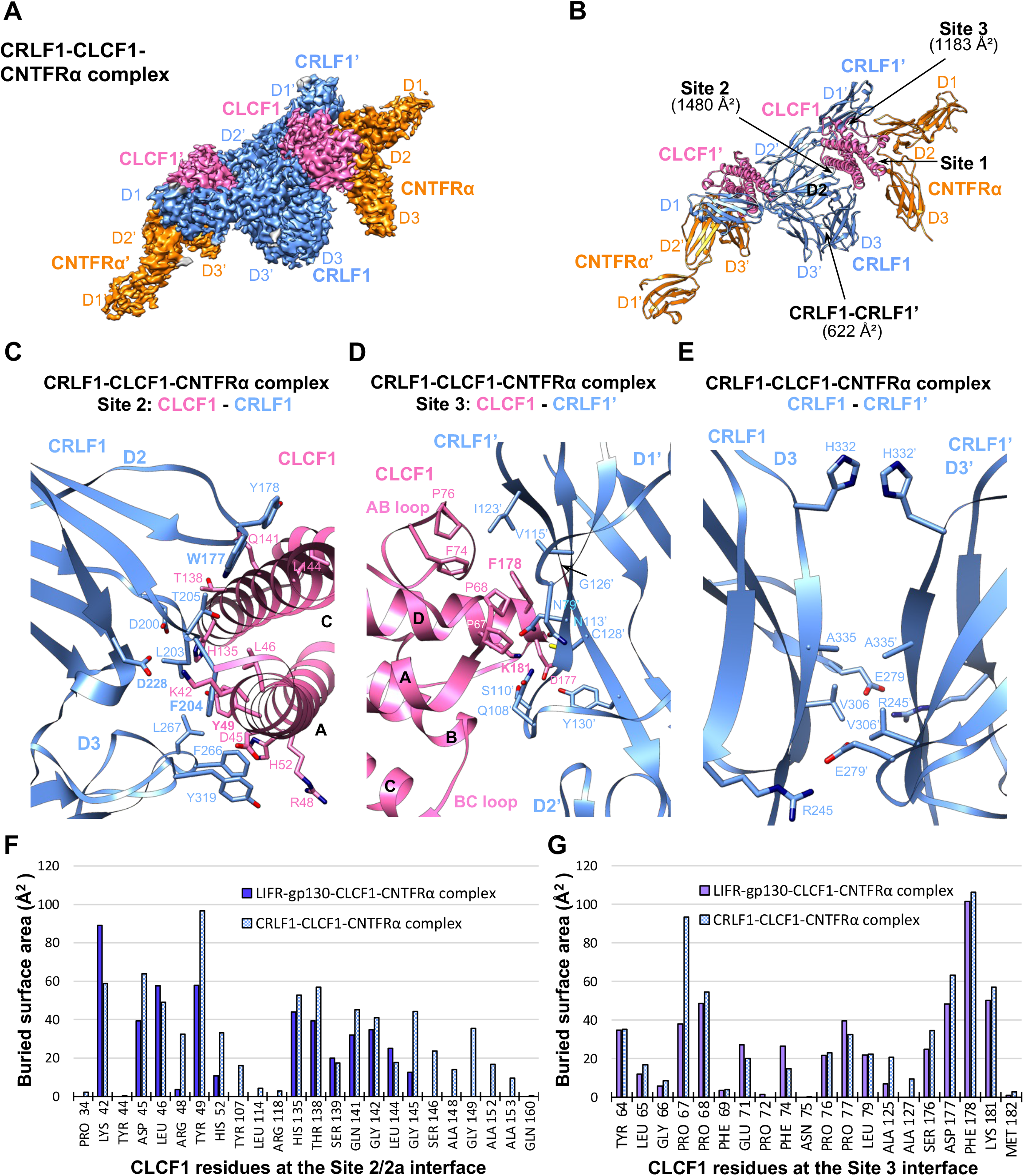
Cryo-EM structure of the CRLF1-CLCF1-CNTFRα complex. (A) Cryo-EM density map of the CRLF1-CLCF1-CNTFRα complex. The two sets of molecules in the hexameric complex are annotated as CRLF1, CLCF1, CNTFRα, CRLF1’, CLCF1’, and CNTFRα’. (B) Cartoon representation of the CRLF1-CLCF1-CNTFRα complex. The Site 2 and 3 interfaces, and CRLF1 dimer interface with corresponding buried surface areas calculated by PDBePISA are indicated by arrows. (C-E) Details of binding interfaces at Site 2, Site 3 and CRLF1 dimer interface as indicated in (B). (F-G) Histograms of buried surface area for CLCF1 residues at Site 2/2a (F) and Site 3 (G) interfaces in the LIFR-gp130-CLCF1-CNTFRα complex (CLCF1 signaling complex) and CRLF1-CLCF1-CNTFRα complex.

The elbow region of CRLF1 D2D3 engages CLCF1 Site 2 mimicking the interactions between gp130 D2D3 and CLCF1 (Fig. 3C). CRLF1 contacts a total of 24 CLCF1 residues, including all 13 residues that are covered by gp130 (Fig. 3F). Therefore, the CRLF1-CLCF1 interface has a much larger buried surface area (1,480 Å^2^) than the gp130-CLCF1 interface (944 Å^2^). Notably, the three gp130 residues crucial for engaging CLCF1, W164, F191, and D215, all have equivalents in CRLF1 at similar locations, i.e., W177, F204, and D228, respectively. These CRLF1 residues interact with CLCF1 analogous to the corresponding gp130 residues. While F204 is inserted into a cavity formed by CLCF1 K42, D45, L46, and Y49 at helix A, W177 packs against helix C and D228 pairs with K42 of CLCF1. The involvement of several other aromatic residues of CRLF1 in the interactions, including Y178, F266, and Y319, further stabilizes the binding. Unlike gp130 D3 which only contributes 9.8% of the buried surface area to the gp130-CLCF1 interface (Fig. 2F), CRLF1 D3 contributes a much larger fraction of buried surface area (24.5%) to the Site 2 CRLF1-CLCF1 interface (Fig. 3C).

Furthermore, the way LIFR D3 captures CLCF1 Site 3 is copied by D1 of another CRLF1 molecule, CRLF1’, in the CRLF1-CLCF1-CNTFRα complex (Fig. 3D). The F*XX*K motif of CLCF1 also dominates the binding to CRLF1’. CLCF1 F178 is stacked against CRLF1’ G126’ and is surrounded by N113’, V115’, I123’, and C128’ of CRLF1’, while CLCF1 K181 coordinates with S110’, N113’, and C128’ of CRLF1’ to form hydrogen bonds. Additionally, key CLCF1 Site 3 residues are all shared by CRLF1’ and LIFR for binding to this site (Fig. 3G). Consistent with our structure, the critical roles of CLCF1 F178 and K181 in engaging CRLF1’ have been supported by alanine substitutions (*18*).

The two CRLF1 molecules also directly contact each other at their D3 domains (Fig. 3E). While R245 of one CRLF1 couples to E279 of another CRLF1 to form hydrogen bonds and salt bridges, H332, A335, and V306 from one molecule pack against the same corresponding residues from another molecule to hold both CRLF1 D3 domains together. However, this interface has a small buried surface area (622 Å^2^) with limited number of residues involved in the interactions.

### Structural characterization of the IL-27 and IL-6 signaling complexes

In order to make a comprehensive analysis for the assembly of the gp130 family cytokine-receptor complexes, we further determined cryo-EM structures of the human IL-27 and IL-6 signaling complexes, which share similar “Sites 1-3” interactions with Site 3 of both ligands engaging gp130 (Figs. 4 and 5).

**Fig. 4.**
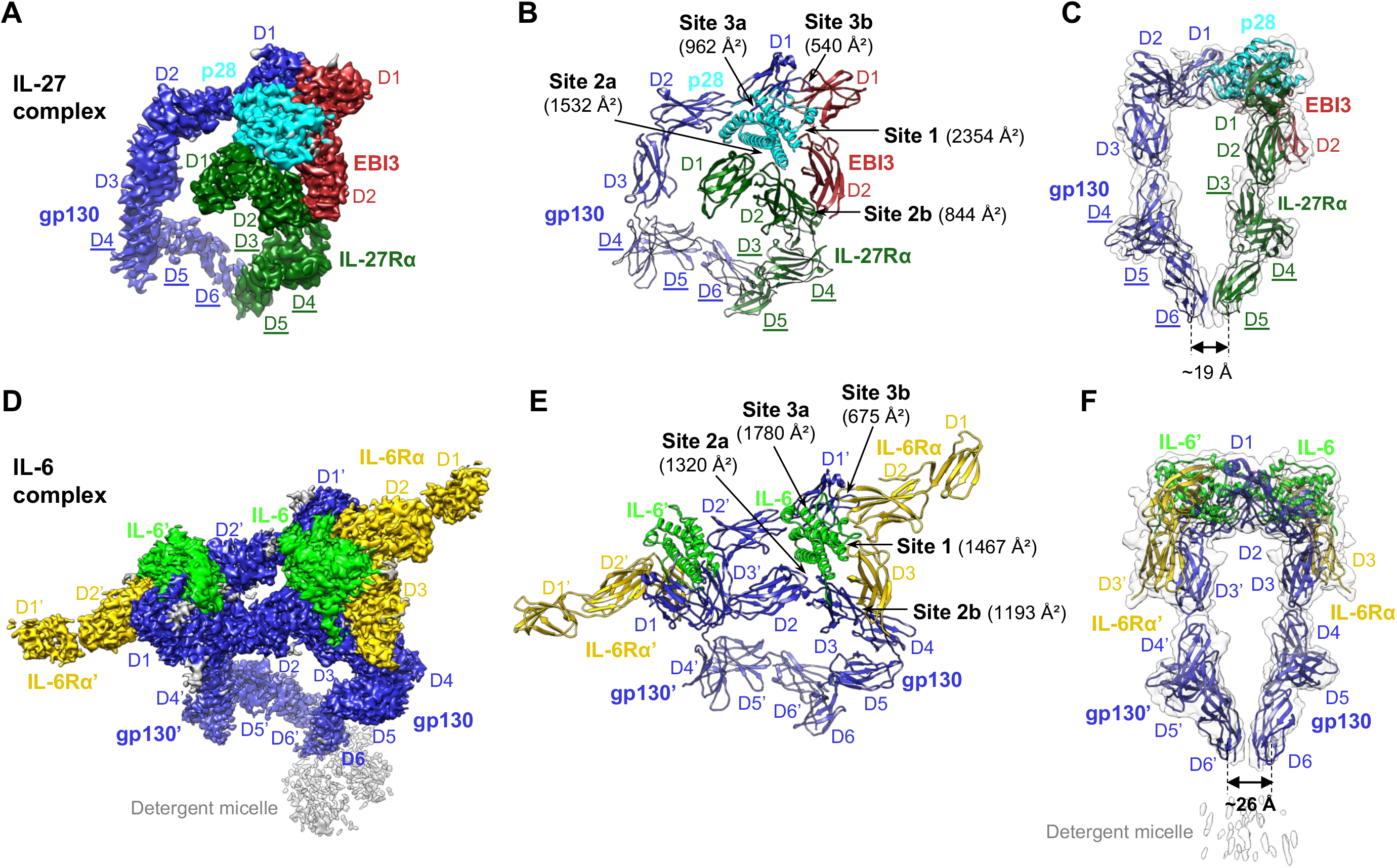
Cryo-EM structures of the IL-27 complex and detergent-solubilized IL-6 complex. (A-C) The IL-27 signaling complex is shown as colored cryo-EM density map (A), ribbon representation of the model (B), and side view of the model in transparent low-pass filtered density map (C). Receptor domains that were only rigid-body refined against the density are underlined. (D-F) Cryo-EM density map (D), cartoon representation (E), and side view of the model in transparent low-pass filtered density map (F) of the IL-6 complex in detergent. The two sets of molecules in the hexameric complex are annotated as gp130, IL-6, IL-6Rα, gp130’, IL-6’, and IL-6Rα’. The interaction interfaces with corresponding buried surface areas calculated by PDBePISA are indicated by arrows in panels B and E. The distance between the bottom centers of the receptor juxtamembrane domains in each complex is estimated and shown in panels C and F.

**Fig. 5.**
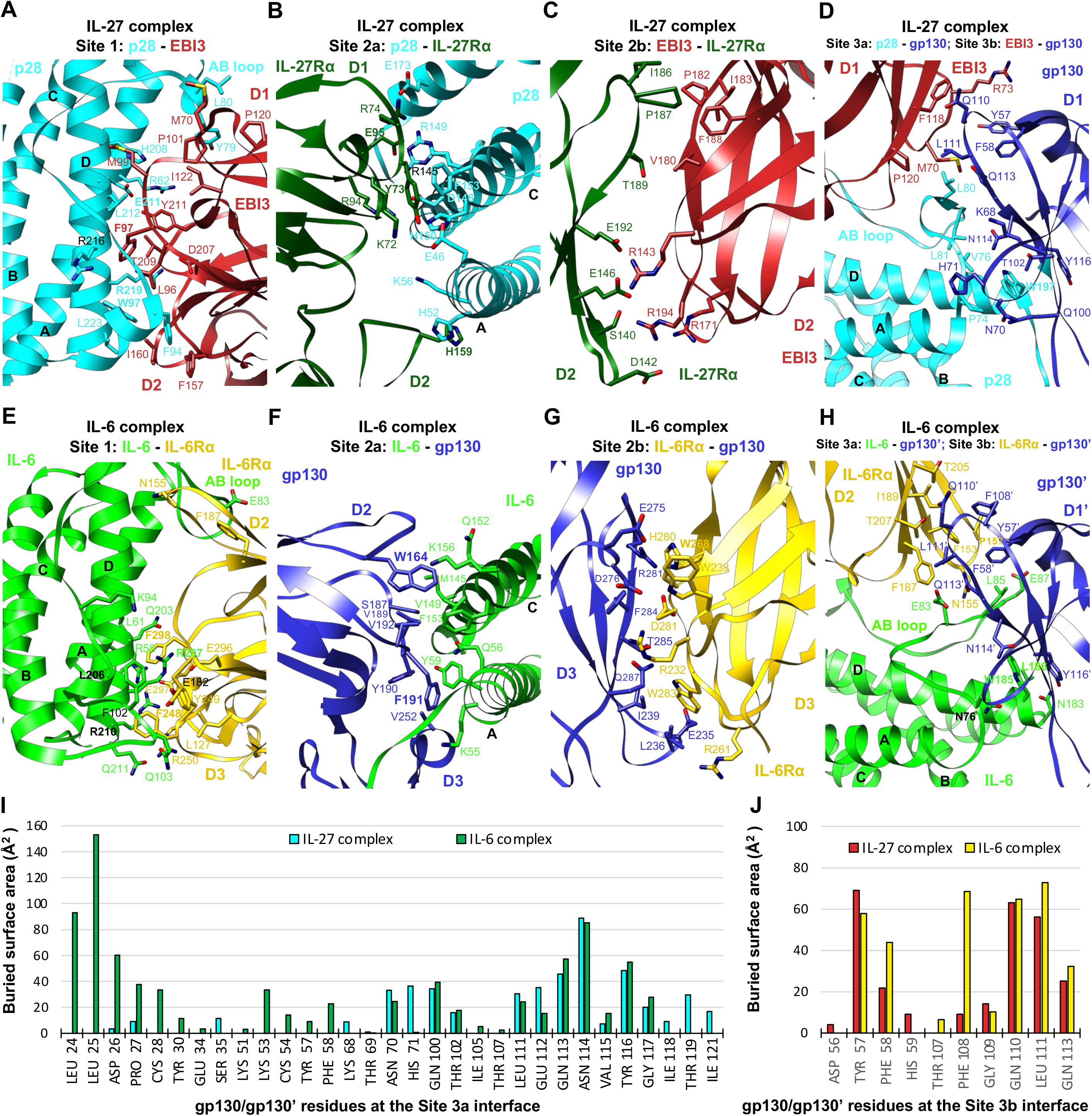
Interaction interfaces of the IL-27 complex and IL-6 complex. (A-D) Details of binding interfaces of the IL-27 complex at Site 1, Site 2a, Site2b and Site 3a/3b as indicated in Fig. 4B. (E-H) Details of binding interfaces of the IL-6 complex at Site 1, Site 2a, Site2b and Site 3a/3b as indicated in Fig. 4E. (I-J) Histograms of buried surface area for gp130/gp130’ residues at Site 3a (I) and Site 3b (J) interfaces in the IL-27 complex and IL-6 complex.

### Cryo-EM structure of the IL-27 signaling complex

IL-27, a heterodimeric cytokine composed of p28 and a soluble receptor EBI3, signals via IL-27Rα and gp130. Unlike CNTF, CLCF1 and LIF which all bind to gp130 D2D3 at Site 2, p28 engages gp130 D1 at Site 3. Using full ectodomains of gp130 and IL-27Rα, and an EBI3-p28 fusion protein linked by a flexible linker, we obtained a 4.14 Å cryo-EM map of the IL-27 quaternary signaling complex that has well resolved density for domain placement and rigid-body refinement (Fig. 4A; fig. S5, A to E). We further ran a focused refinement of the interaction core region, including p28, EBI3, IL-27Rα D1D2, and gp130 D1-D3, and obtained a 3.81 Å map with ~3.40 Å local resolution around the interaction interfaces, which was used for model building (fig. S5, F to H). A published structure of gp130 D4-D6 (*2*) and a mode of IL-27Rα D3-D5 predicted by AlphaFold were used to generate a more complete model for the IL-27 complex (Fig. 4B). Intriguingly, an acute bend analogous to that found in gp130 and LIFR, also exists between D3 and D4 of IL-27Rα. This bend, together with the bend of gp130 at D4D5, brings the receptor juxtamembrane domains into close proximity (~19 Å between the bottom centers of gp130 D6 and IL-27Rα D5) (Fig. 4C).

EBI3 makes a broad contact with p28 helices A and D and the AB loop at the Site 1 interface to bury a surface area of up to 2,354 Å^2^ (Figs. 4B and 5A). A network of hydrogen bonds and salt bridges mediate the interactions, with extensive hydrophobic interactions further stabilizing the binding. The interface is centered around E211 and R219 of p28, which form hydrogen bonds with Y211 and T209 of EBI3, respectively. EBI3 F97, similar to the evolutionarily conserved CNTFRα residue F172 in the CNTF and CLCF1 complexes, contributes a large buried surface area (152 Å^2^) in the interface. Y79 and L80 at the N-terminus of the p28 AB loop from a hydrophobic patch with M70, P101, P120, and I122 of EBI3. Another hydrophobic patch is formed at the C-terminus of this loop, with F94, W97, and L223 of p28 engaging L96, F157, and I160 of EBI3. Consistent with our data, previous mutagenesis work has shown that EBI3^F97^ and p28^W97^ are both critical for IL-27 signaling (*24*). Notably, the equivalent residues of p28^W97^ at similar locations of CNTF and CLCF1, CNTF^W64^ and CLCF1^W94^, are both crucial for interacting with CNTFRα (Fig. 2, A and E).

The p28 ligand binds to the hinge of IL-27Rα CHR (D1D2) at the Site 2a interface (Fig. 5B), which is dominated by hydrogen bond and salt bridge interactions mediated by charged residues, including K72, R74, R94 and E95 of IL-27Rα and E46, K56, R145, D146, and E173 of p28. In addition, Y73 in D1 and H159 in D2 of IL-27Rα pack against p28^H150, F153^ at the C-terminus of helix C, and p28^K56, H52^ at the middle of helix A, respectively. Notably, of the 18 IL-27Rα residues contacting p28, Y73, E95, and H159 make up ~40% of the buried surface area on the IL-27Rα side.

The “sandwiched” position of p28 between EBI3 and IL-27Rα is stabilized by the Site 2b interactions mediated by IL-27Rα D2 and EBI3 D2 (Fig. 5C). This interface is enriched with negatively charged residues on IL-27Rα (D142, E146, and E192) and positively charged residues on EBI3 (R143, R171, and R194), which mediate electrostatic interactions. A hydrophobic patch of residues, IL-27Rα^I186, P187^ and EBI3^P182, I183, F188^, further improves the binding. The Site 2b interface adds an additional 844 Å^2^ buried surface area to Site 2a, making a total buried surface area of 2,376 Å^2^ for the composite Site 2 and leading to increased binding interactions, which might explain why p28 activity is EBI3-dependent (*25*).

Similar to the IL-6 complex (*3*), the IL-27 complex has a composite Site 3 interface, including Site 3a formed by gp130 Ig domain (D1) engaging p28, and Site 3b formed by gp130 D1 contacting EBI3 D1 (Figs. 4B and 5D). Site 3a has a relatively limited buried surface area (962 Å^2^) with W197 at the N-terminus of p28 helix D packing against gp130^Y116^ to serve as a binding anchor. The aromatic residue W197, which is equivalent to a Phenylalanine residue in the F*XX*K motif of CNTF (F152), CLCF1 (F178), and LIF (F178), is evolutionarily conserved in IL-6 (W185) (*3*) and vIL-6 (W166) (*23*), and has been shown to be crucial for IL-27 signaling (*25*). A group of residues at the N-terminus of p28 AB loop (V76, L80, L81) make additional hydrophobic contacts with gp130 D1. The Site 3b interface is centered around EBI3^F118^ and has a small buried surface area (540 Å^2^). The tip of gp130 D1 leans against the top side of EBI3 D1 and the interactions are largely hydrophobic.

### Cryo-EM structure of the IL-6 signaling complex in detergent

We further characterized the IL-6 signaling complex in which the ligand binds to gp130 at both Site 2 and Site 3. With the goal of obtaining structural information for the TM domain, we purified gp130 with the TM region and cytoplasmic Box1/Box2 motifs and reconstituted the complex with IL-6Rα ectodomain and IL-6 in detergent. We obtained a 3.22 Å cryo-EM map of this hexameric complex with 2-fold symmetry (Fig. 4, D and E; fig. S6). Although a density corresponding to the detergent micelle is present, the TM helix and cytoplasmic region of gp130 are not resolved, and there is a ~15 Å gap between gp130 C-terminal density and the detergent micelle (fig. S6C), indicating some level of flexibility around gp130 TM domain even though the two TM helices are embedded inside the detergent micelle. The acute bend of gp130 at D4D5 brings the bottom centers of the two gp130 juxtamembrane domains to ~26 Å apart (Fig. 4F).

The assembly of the IL-6 complex interaction core region agrees with the 3.65 Å crystal structure of gp130 D1-D3/IL-6Rα D2D3/IL-6 complex (*3*). Briefly, the Site 1 interface is dominated by IL-6Rα^F248, F298^ and IL-6^R207, R210^ through a network of hydrophobic and electrostatic interactions. The binding center at IL-6 Site 1 is close to the C-terminus of helix D with IL-6Rα D3 contributing the majority (63.7%) of the buried surface area (Fig. 5E). In contrast, the Site 1 binding centers of CNTF, CLCF1, and IL-27 p28 are all close to the middle of helix D with the C-terminal domains of the corresponding alpha receptor (CNTFRα D3, EBI3 D2) contributing less to the binding than the preceding domains (Fig. 2, A and E; Fig. 5A). At Site 2a of the IL-6 complex, W164 and F191 of gp130 engage IL-6 helices A and C, respectively in a similar manner as they interact with CNTF, CLCF1, and LIF (Fig. 5F). The Site 2b interface stabilizes the composite Site 2 binding by introducing extra buried surface area with gp130^E235, D276, R281^ and IL-6Rα^R232, D281, W283^ playing key roles (Fig. 5G). The Site 3a interface is characterized by W185 on the C-terminus of IL-6 helix D stacking with Y116’ on D1’ of the second gp130 receptor, gp130’ (Fig. 5H). Notably, the long AB loop of IL-6 is inserted into a broad cavity on the head of gp130’ D1’ and the N-terminal end of gp130’ extends in parallel with one side of the loop to mediate extensive interactions, both of which are not seen in the IL-27 complex (Fig. 5, D, H, and I). These differences lead to a much larger buried surface area at Site 3a of the IL-6 complex (1,780 Å^2^) than that of the IL-27 complex (962 Å^2^). Finally, the Site 3b interface of the IL-6 complex is made by the tip of gp130’ contacting the side of IL-6Rα D2 and is centered around IL-6Rα^F153^ (Fig. 5H). Key gp130’ residues that contact IL-6Rα at this interface also engage EBI3 in the IL-27 complex (Fig. 5J).

### Structural similarities of cytokines and non-signaling receptors in the gp130 family cytokine-receptor complexes

We next compared six gp130 family cytokines, including five characterized in this study and OSM which has a published structure (*26*). Despite the low sequence identities of these cytokines (table S1), they all share a canonical four-helix bundle structure, where four major helices are linked by three loops (Fig. 6A). Notably, some key residues for receptor binding at Site 1 and Site 3 are conserved across different human cytokines, including CNTF^W64^, CLCF1^W94^, and IL-27 p28^W97^ at Site 1, CNTF^F152, K155^, CLCF1^F178, K181^, LIF^F178, K181^, and OSM^F185, K188^ at Site 3 that engages LIFR, as well as IL-27 p28^W197^ and IL-6^W185^ at Site 3 that binds to gp130. Of the five gp130 family cytokine signaling complexes we examined, the LIF complex is the only one without a non-signaling alpha receptor. Notably, the N-terminal extension of LIF preceding helix A is tethered to helix C by two disulfide bonds (C34-C156 and C40-C153) so as to adopt a rigid conformation to make extensive contacts with gp130 D2 (Fig. 2J). The additional buried surface area introduced by these contacts might serve to improve LIF Site 2 affinity to gp130, which might explain why LIF does not require an alpha receptor for signaling. Another gp130 family cytokine, OSM, shares this alpha receptor independence to signal though gp130 and either LIFR or OSM receptor. Interestingly, OSM also has an N-terminal extension that is tethered to helix C by the C31-C152 disulfide bond (*26*) and might also increase OSM Site 2 affinity to gp130. In contrast, a similar conformation of a rigid N-terminal extension tethered to helix C by disulfide bonds is not observed in CNTF, CLCF1, IL-27 p28, or IL-6, which all require a non-signaling receptor for signal transduction. Similar to the N-terminal extension of LIF which makes additional contacts with gp130 D2 to increase the buried Site 2 surface area, the non-signaling receptors in the CNTF, CLCF1, IL-6, and IL-27 complexes all serve to increase the total buried surface area of the composite Site 2 in these complexes by making extra Site 2b contacts with the corresponding signaling receptor bound to Site 2a (Figs. 1 and 4).

**Fig. 6.**
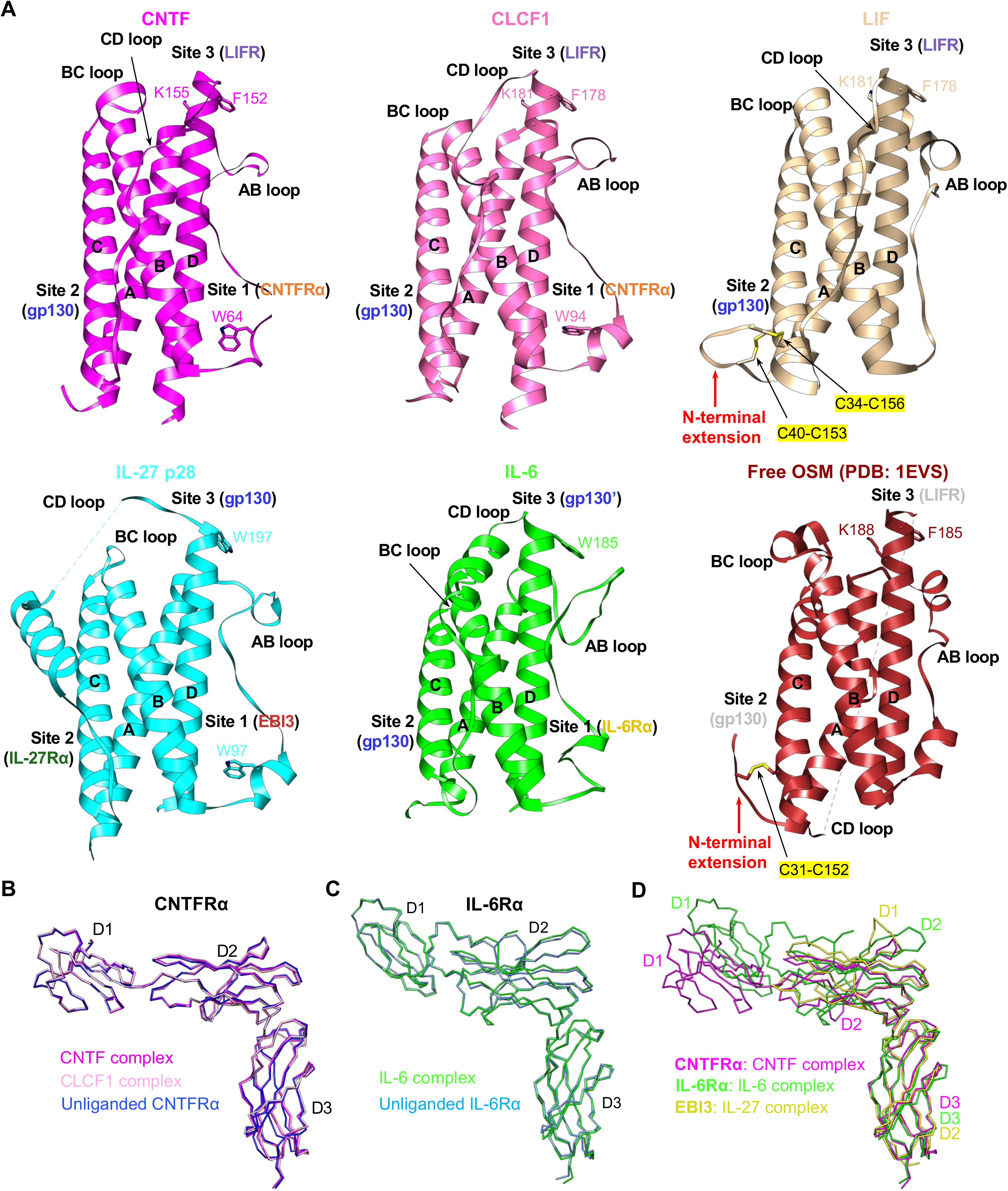
Structural comparisons of cytokines and non-signaling receptors in the gp130 family cytokine-receptor complexes. (A) The structures of six gp130 family cytokines are aligned on the four-helix bundle. Important conserved residues for binding to receptors at Site 1 and Site 3 are shown in stick form and labeled. LIF and OSM both have a N-terminal extension preceding helix A that is tethered to helix C by disulfide bonds (C34-C156 and C40-C153 for LIF; C31-C152 for OSM), which is not seen in other cytokines. (B) Superposition of CNTFRα from the CNTF and CLCF1 signaling complexes, as well as unliganded CNTFRα from the CNTFRα/REGN8938 Fab/H4H25322P2 Fab complex. All molecules are shown as C-alpha ribbon traces. (C) Superposition of IL-6Rα from the IL-6 signaling complex and unliganded IL-6Rα (PDB: 1N26) (D) Superposition of liganded CNTFRα, IL-6Rα, and EBI3 from the CNTF, IL-6, and IL-27 signaling complexes, respectively.

We further compared the three non-signaling receptors characterized in our study, including CNTFRα, IL-6Rα, and EBI3. No significant conformational changes of CNTFRα and IL-6Rα were observed upon ligand binding (Fig. 6, B and C). Notably, CNTFRα, IL-6Rα, and EBI3 adopt highly similar conformations at the elbow regions where the ligands bind (Fig. 6D), which may explain why IL-6Rα could also serve as an alpha-receptor for CNTF and IL-27 p28 instead of CNTFRα and EBI3, respectively (*27*, *28*).

## Discussion

It has been very challenging to characterize the structures of the ‘tall’ gp130 family cytokine receptors due to their common elongated geometry and resulting flexibility. To our knowledge, this paper represents the first high-resolution cryo-EM structure determination of the full ectodomains of this family of receptors, including gp130, LIFR, and IL-27Rα. Surprisingly, in the complexes we characterized, gp130 and LIFR are both quite rigid overall, despite some degree of flexibility at the membrane-proximal regions (Fig. 7, A and B). There are no large conformational changes of the receptors upon cytokine binding as well. gp130 CHR (D2D3) engages CNTF, CLCF1, LIF and IL-6 at Site 2 in a highly similar manner, with 10 gp130 residues (W164, T166, H167, S187, V189, Y190, F191, V192, N193, V252) being shared by all four cytokines and F191 playing a key role for binding (Fig. 7C). The ways in which LIFR engages CNTF, CLCF1 and LIF are even more conserved, with 17 LIFR residues being shared by the three cytokines (Fig. 7D). Important LIFR residues for binding include S310, N313, I322, and V326.

**Fig. 7.**
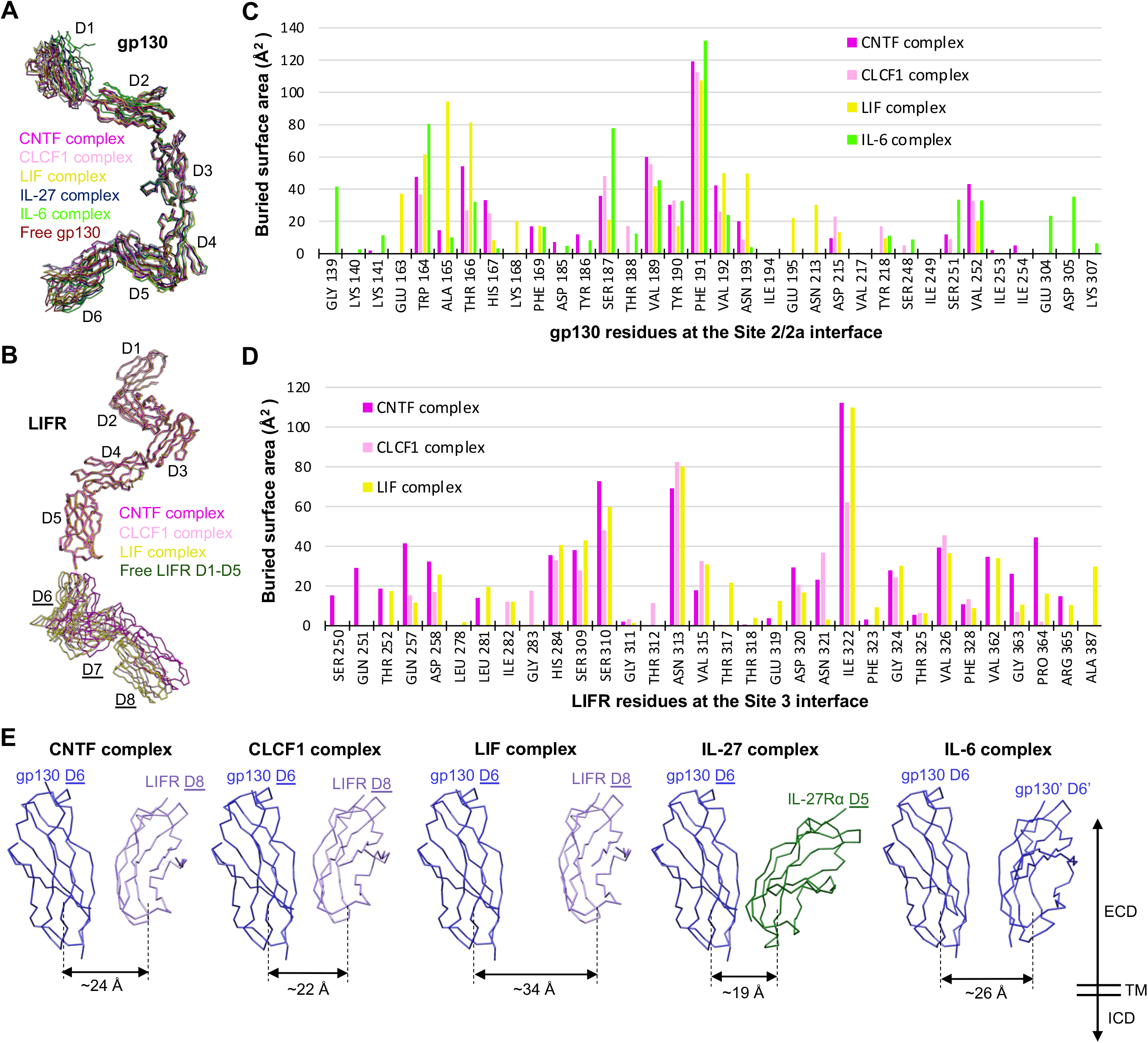
Conformations of shared signaling receptors and relative positions of the receptor juxtamembrane domains in different gp130 family cytokine signaling complexes. (A) Superposition of gp130 from the CNTF, CLCF1, LIF, IL-27, and IL-6 complexes, as well as free gp130 (PDB: 3L5H). All molecules are shown as C-alpha ribbon traces. (B) Superposition of LIFR from the CNTF, CLCF1, and LIF complexes, as well as free LIFR D1-D5 (PDB: 3E0G). (C) Histogram of buried surface area for gp130 residues at the Site 2/2a interface in various signaling complexes. (D) Histogram of buried surface area for LIFR residues at the Site 3 interface in various signaling complexes. (E) Relative positions of membrane-proximal domains of the two signaling receptors in different gp130 cytokine signaling complexes. All models are aligned on gp130 D6. The approximate distance between the bottom centers of the receptor juxtamembrane domains in each complex is indicated. ECD: extracellular domain; TM: transmembrane domain; ICD: intracellular domain.

A common feature of the ‘tall’ signaling receptors, gp130, LIFR, and IL-27Rα, is an acute bend (~80°) between the first and second FNIII domains (i.e., gp130 D4D5, LIFR D6D7, IL-27Rα D3D4), which is a crucial geometry for signaling. The bends at the two signaling receptors in each of the signaling complexes we examined serve to bring the bottom centers of the receptor juxtamembrane domains to around 30 Å. This is similar to the distances between the receptor juxtamembrane domains observed in multiple other cytokine-receptor complexes that also activate the JAK/STAT pathway (fig. S7), including the Epo-EpoR complex (*29*), insulin-insulin receptor complex (*30*), and IGF1-IGF1R complex (*31*). Moreover, the ~30 Å distance is also comparable to the distance between the two membrane-proximal FERM-SH2 domains of dimeric JAK1 bound to Box1/Box2 motifs of a cytokine receptor on the intracellular side (*32*). These consistent observations suggest that bringing the two signaling receptor juxtamembrane domains to ~30 Å apart might be a prerequisite for activating the JAK/STAT pathway. The EM density of these juxtamembrane domains in most of our maps does not permit detailed analysis of residue-residue interactions; however, we note that the modeled positions of the domains with the closest approach (e.g., in the IL-27 and CLCF1 complexes) will place residues in those domains close enough for direct contact, with Cα-Cα distances <6 Å. The two gp130 D6 domains in the IL-6 complex are better resolved, likely because the insertion of TM helices into the detergent micelle has restrained the flexibility of these domains. Consistent with the observation that no D6-D6 contacts are made in gp130 crystal lattice (*2*), we do not see direct interactions between the gp130 juxtamembrane domains in the IL-6 signaling complex. Furthermore, the successful reconstitution of the p28/EBI3/gp130 D1-D3/IL-27Rα D1D2 complex (*13*), which has C-terminal truncations of both signaling receptors, indicates that even if gp130 D6 directly contacts IL-27Rα D5 in the full IL-27 signaling complex as it appears in the EM map (Fig. 4C), this juxtamembrane interaction is not essential for the assembly of the complex.

The gp130 family cytokines all activate the JAK/STAT pathway, but how the signaling specificity is derived remains unclear. For example, it is not clear why IL-27 activates both STAT1 and STAT3, while IL-6 predominantly signals via STAT3 phosphorylation (*33*). Our data show that these cytokines do have variable loop conformations, especially at the N-terminus of the AB loop that makes up part of the Site 3 receptor binding epitope (Fig. 6A), which could lead to distinct binding topologies of Site 3 receptors. Moreover, even though the Site 2 receptor binding epitope is located on helix A and C which are more structurally conserved across the cytokines, different binding modes could still be adopted by the shared receptor for different cytokines. For example, although Site 2 of CNTF, CLCF1, LIF and IL-6 all engage gp130, the binding location of gp130 on LIF is apparently shifted to the N-terminal end of helix A compared to that of other cytokines, leading to an enlarged distance between the membrane-proximal domains of the two signaling receptors in the LIF complex. These different topologies of binding to various cytokines, together with divergent ways of engaging alpha receptors, lead to distinct angles and distances of the receptor juxtamembrane domains in different complexes (Fig. 7E). It is possible that these ectodomain topology differences of signaling receptors could be transmitted into the intracellular domains (ICDs), which in turn affect the orientation or proximity of the JAK kinases bound to the ICDs of the receptors. As a result, this may alter JAK-mediated phosphorylation events on various STAT substrates and adaptors, and lead to distinct gene expression profiles. In agreement with this hypothesis, engineered Epo-EpoR signaling complexes with different orientations or distances of receptor juxtamembrane domains induce distinct effects in hematopoiesis (*34*).

Difference in receptor-cytokine affinity and complex stability could be another factor that affects the biological specificities of the cytokines. Engineered IL-6 with lower affinity to gp130 was shown to decrease STAT1 phosphorylation more profoundly than STAT3 phosphorylation and thereby induce STAT3 biased responses (*35*). IL-13 variants with various affinities to receptors also cause different functional outputs (*36*). Both studies proposed that changes in cytokine-receptor affinities alter stability of the signaling complex, which in turn affects receptor endocytosis that plays an important role in regulating STATs activation pattern. We found that the CLCF1 complex and IL-27 complex appear to be much more unstable than other complexes on the EM grid, since both samples contained an extremely low percentage of full complex particles (~2%). We also observed various assembly intermediates in the CNTF complex (CNTF-CNTFRα-gp130 intermediate) (fig. S1C), LIF complex (LIF-LIFR intermediate) (fig. S3H), and IL-27 complex (p28-EBI3-IL-27Rα intermediate) (fig. S5C), likely suggesting different relative binding affinities of the Site 2 and Site 3 receptors in different complexes. These differences in affinity and stability of the cytokine-receptor complexes could also possibly contribute to biological specificities of the gp130 family cytokines in cells as previously proposed (*35*).

CRLF1 is known to be critical for CLCF1 secretion but its role in CLCF1 signaling remains elusive. It was shown that the two key CLCF1 residues (F178 and K181) engaging LIFR are also important for binding to CRLF1, suggesting that CRLF1 and LIFR compete for binding to CLCF1 Site 3 and thereby CLCF1 may need to be released from CRLF1 for signaling (*18*). Another study reported that CRLF1 is able to form a tripartite complex with CLCF1 and CNTFRα and promote CLCF1 signaling by sustaining CLCF1 binding to CNTFRα (*22*). However, our structure of the CRLF1-CLCF1-CNTFRα complex shows that CLRF1 does not contact CNTFRα directly and CNTFRα is not required for CLCF1-meditated CRLF1 dimerization. Surprisingly, CLCF1 Site 2 and Site 3 are engaged by two CRLF1 molecules analogous to how they are engaged by gp130 and LIFR in the CLCF1 signaling complex, supporting the model that CLCF1 has to dissociate from CRLF1 in order to bind to gp130 and LIFR to form a functional signaling complex (*18*). The mechanism for CLCF1 release from CRLF1 in cells remains to be investigated.

Our structures of the gp130 family cytokine-receptor complexes expand our view of the signaling mechanism of this family and provide valuable insights for therapeutically targeting gp130-mediated signaling pathways. Based on the detailed cytokine-receptor contacts described above, it may be possible to generate chimeric cytokines, agonists, or antagonists via structure-guided protein engineering. The geometry of the full extracellular portion of each signaling complex could also guide the design of therapeutics such as bispecific antibodies to bring the signaling receptors together in a conformation that mimics the natural cytokine signaling complex. Engineered molecules such as these will hopefully be valuable in treating diseases arising from disorders in gp130-mediated signaling.

## Acknowledgments

The authors would like to thank John C. Lin, Matthew Sleeman, Erich Goebel and Luke McGoldrick for critical reading of the manuscript, Jean Yanolatos for project management, Judith Altarejos for discussion of the project, Yang Shen for early discussion on model building of the CNTF signaling complex, and Olav Olsen for help with the submission of the manuscript.

## Funding

This project has been funded by Regeneron Pharmaceuticals.

## Author contributions

Y.Z., P.E.S., G.D.Y., A.J.M., M.W.S., W.C.O., and M.C.F. conceptualized the studies. Y.Z. prepared complexes, acquired cryo-EM data, processed data, and built and refined the atomic models, with contributions from K.S. and M.C.F.. P.E.S and J.C. contributed most of the reagents. J.W., A.G.Z, and S.H. conceptualized the IL-27 signaling complex study and analyzed the IL-27 complex structural data. G.D.Y., A.J.M., M.W.S., W.C.O., and M.C.F. analyzed data and supervised the whole project. Y.Z. drafted the manuscript, which was edited and finalized with contributions of all authors.

## Competing interests

Regeneron authors own options and/or stock of the company. The two anti-CNTFRα antibodies used in this study have been described in a pending provisional patent application. G.D.Y., A.J.M., M.W.S., and W.C.O. are officers of Regeneron.

## Data and materials availability

Regeneron materials described in this manuscript may be made available to qualified, academic, noncommercial researchers through a materials transfer agreement upon request at https://regeneron.envisionpharma.com/vt_regeneron/. For questions about how Regeneron shares materials, use the email address. Protein Data Bank (PDB) and Electron Microscopy Data Bank (EMDB) identification numbers for the complexes described in this paper were listed in tables S2.

