## Supplementary Material for "Structural insights into the assembly of gp130 family cytokine signaling complexes"

### Materials and Methods

#### Protein purification

All proteins used in this study were recombinant human proteins. The protein residues were numbered based on their UniProt sequences; the signal peptides were counted in the numbering. The following proteins were all C-terminal myc-myc-His (mmH)-tagged secreted proteins expressed in Chinese Hamster Ovary (CHO-K1) cells: gp130 ectodomain (aa E23-E619, REGN2669), LIFR ectodomain (aa Q45-S833, REGN3269), CNTFRα (aa Q23-S342)-GGGPG-CNTF (aa M1-I186) fusion protein (REGN3637), CNTFRα (aa Q23-S342, REGN3000), IL-27Rα ectodomain (aa Q33-K516, REGN9497), EBI3 (aa R21-K229)-$(GGGGS)_4$-p28 (aa F29-P243) fusion protein (REGN5948), and IL-6Rα ectodomain (aa L20-M331, REGN78). Filtered cell culture supernatants containing target proteins were buffer exchanged through dialysis against DPBS and loaded onto pre-equilibrated Talon columns (Clontech, #635682). After washing the columns with DPBS containing 500 mM NaCl, followed by a second wash with DPBS plus 500 mM NaCl and 5 mM Imidazole, the proteins were eluted with DPBS plus 500 mM NaCl and 200 mM Imidazole. The eluates were dialyzed against DPBS with 5% glycerol, and subsequently, the proteins were further purified by size exclusion chromatography (SEC). The proteins were concentrated and frozen for future use. IL-6 (aa V30-M212, REGN125) was expressed as an inclusion body in E.coli and refolded into a soluble form.

Human gp130 with full ectodomain, transmembrane domain and cytoplasmic Box1/2 region linked to a hFc tag ((gp130 E23-D700)-$(GGGGS)_3$-hFc)) was expressed in the Expi293 expression system (ThermoFisher). The cell pellet was homogenized in lysis buffer (PBS + 1% DDM detergent). Clarified lysate was loaded onto hand packed MabSelect SuRe (Cytiva) column. The column was washed with lysis buffer and then PBS with 0.02% DDM (Anatrace). Bound gp130 protein was eluted four times with batches of 2 column volume elution buffer containing 100 mM glycine pH 2.7, 150 mM NaCl, 0.02% DDM into tubes containing 1 ml 1M Tris-Cl pH 8.0 for neutralization. Fractions were analyzed by SDS-PAGE, and gp130-containing fractions were combined and dialyzed against buffer containing 20 mM HEPES pH 7.4, 150 mM NaCl, 5% glycerol, and 0.02% DDM. The protein was further concentrated with a 100 kDa MWCO centrifugal concentrator and flash frozen in liquid nitrogen for future use.

Three other proteins were purchased from R&D Systems, including CNTFRα (aa Q23-P346)-GSGSSRGGSGSGGSGGGGSKL-CLCF1 (L28-F225) fusion protein (#2415-CR) and CRLF1 (aa A38-R422)/CLCF1 (aa L28-F225) complex protein (#1151-CL) purified from Mouse myeloma cell line NS0, and LIF (aa P24-F202, #7734-LF) purified from E.coli.

#### Fab fragment preparation

Two anti-CNTFRα antibodies, REGN8938 and H4H25311P2, were digested into $F(ab')^2$ and Fc fragments using Fabricator enzyme (Genovis) following instructions from the manufacturer. $F(ab')^2$ was reduced into F(ab)' using 2-mercaptoethylamine (2-MEA, ThermoFisher) followed by Fc fragment removal using the CaptureSelect IgG-Fc (ms) affinity resin (ThermoFisher). F(ab)' fragments were further purified over a Superdex 200 increase 10/300 GL gel filtration column equilibrated with 50 mM Tris pH 7.5, 150 mM NaCl, and concentrated using a 10 kDa MWCO centrifugal concentrator.

**Complex preparation**
All complexes were reconstituted by mixing the corresponding components at an equal molar ratio followed by incubation at 4 °C for 1 h as follows: CNTF complex (gp130 ectodomain, LIFR ectodomain, and CNTFRα-CNTF fusion protein), CNTFRα/REGN8938 Fab/H4H25311P2 Fab complex (CNTFRα, REGN8938 Fab, and H4H25311P2 Fab), CLCF1 complex (gp130 ectodomain, LIFR ectodomain, and CNTFRα-CLCF1 fusion protein), CRLF1/CLCF1/CNTFRα complex (CRLF1/CLCF1 complex and CNTFRα), LIF complex (gp130 ectodomain, LIFR ectodomain, and LIF), IL-27 complex (gp130 ectodomain, IL-27Rα ectodomain, and EBI3-p28 fusion protein), IL-6 complex (gp130 solubilized in detergent, IL-6Rα ectodomain, and IL-6).

All complexes except the IL-6 complex were purified over a Superdex 200 increase 10/300 GL gel filtration column equilibrated with 50 mM Tris pH 7.5, 150 mM NaCl. Peak fractions containing the target complex were collected and concentrated to ~2.5 mg/ml using a 30 kDa MWCO centrifugal concentrator. The IL-6 complex solubilized in detergent was purified over a Superose 6 increase 10/300 GL gel filtration column equilibrated with 50 mM Tris pH 7.5, 150 mM NaCl, and 0.02% GDN (Anatrace), and concentrated with a 100 kDa MWCO centrifugal concentrator to ~1.5 mg/ml.

**Cryo-EM sample preparation and data collection**
Each freshly purified complex was mixed with ~0.15% Amphipol PMAL-C8 (Anatrace) immediately before pipetting 3.5 µL of the mixture onto a UltrAufoil R1.2/1.3, 300 mesh grid (Quantifoil). The grid was blotted for 4 s at a force of 0 and plunge frozen into liquid ethane using a Vitrobot Mark IV (ThermoFisher) operated at 4° C and 100% humidity. The grid was then loaded into a Titan Krios G3i microscope (ThermoFisher) equipped with a K3 camera and energy filter (Gatan) for data collection in counted mode at a nominal magnification of 105,000x using the EPU software (ThermoFisher). Each movie contained 46 dose fractions over a 2 s exposure, and the total acquired dose per Å$^2$ was ~40 electrons. CNTF complex, CNTFRα/REGN8938 Fab/H4H25311P2 Fab complex, and IL-6 complex had a pixel size of 0.85 Å while all other complexes had a pixel size of 0.86 Å. All movies had a defocus range of -1.4 ~ -2.6 µm. The total number of movies collected for each sample was as follows: CNTF complex (6,511), CNTFRα/REGN8938 Fab/H4H25311P2 Fab complex (7,859), CLCF1 complex (25,442), CRLF1/CLCF1/CNTFRα complex (9,943), CRLF1/CLCF1 complex (7,275), LIF complex (11,007), IL-27 complex (26,610), and IL-6 complex (12,143). Cryo-EM data collection, processing and refinement statistics were also summarized in Table S2.

**Cryo-EM data processing**
Cryo-EM data were processed using Cryosparc (*3*). Movies were motion corrected by Patch motion correction and CTF parameters were estimated by Patch CTF estimation. Particles were initially picked using Blob picker to generate 2D class averages to be used as templates for the subsequent template picking. Junk particles were removed by multiple rounds of 2D classification, followed by Ab initio reconstruction, Homogeneous refinement, and Heterogenous refinement to identify the best class of particles representing the target complex. These particles were further refined using non-uniform refinement (*5*) and/or local refinement to generate the final cryo-EM density map.

The number of total particles (a), number of particles subjected to Ab initio reconstruction (b), number of particles used in final refinement (c), and global resolution of the final cryo-EM density map based on a criterion of 0.143 Fourier shell correlation (d) for each sample were as follows: CNTF signaling complex  (a: 3,197,974; b: 778,622; c: 250,735; d: 3.03 Å), CNTFRα/REGN8938 Fab/H4H25311P2 Fab complex (a: 4,515,671; b: 1,046,202; c: 568,328; d: 2.93 Å), CLCF1 signaling complex (a: 13,119,612; b: 202,209; c: 92,463; d: 3.90 Å), CRLF1/CLCF1/CNTFRα complex (a: 1,258,062; b: 203,337; c: 117,773 d: 3.40 Å), CRLF1/CLCF1 complex (a: 1,504984; b: 369,658; c: 204,227; d: 3.45 Å), LIF signaling complex (a: 5,545,456; b: 726,434; c: 171,328; d: 3.54 Å), IL-27 signaling complex (a: 19,480,836; b: 429,793; c: 139,752; d: 4.14 Å for full map and 3.81 Å for focused refinement map), and IL-6 signaling complex (a: 3,881,621; b: 324,482; c: 105,760; d: 3.22 Å).

A single final cryo-EM density map was generated for most complexes. For the CNTF signaling complex, refinement of 250,735 particles led to a 3.03 Å map which has well resolved (~2.8 Å) interaction core region but fragmented LIFR D6-D8 density. This map was used for model building and refinement of the assembly core region. These particles were further subjected to heterogeneous refinement, identifying 100,013 particles with more homogeneous LIFR D6-D8 density. Non-uniform refinement of this subset of particles yielded another map with lower global resolution (3.37 Å) but improved LIFR D6-D8 density. This map was used for fitting and rigid-body refinement of a LIFR D6-D8 structure predicted by AlphaFold (*6*) to generate a more complete model for the CNTF complex that was shown in Figure 1A and 1C. For the IL-27 signaling complex, final refinement of 139,752 particles resulted in a 4.14 Å map which was used for fitting and rigid-body refinement of available gp130 D4-D6 structure (PDB:3L5H) and a model of IL-27Rα D3-D5 predicated by AlphaFold to make a full model for the complex. The interaction core region of the complex, including p28, EBI3, gp130 D1-D3, and IL-27Rα D1-D2, was further subjected to particle subtraction and local refinement, yielding an improved local map with 3.81 Å overall resolution, which was used for building the atomic model of the interaction core region.

**Model building and refinement**

The published structures of gp130 D1-D6 (PDB: 3L5H) (*7*), LIFR D1-D5 (PDB: 3E0G) (*8*), CNTF (PDB: 1CNT) (*9*), CNTFRα D3 (PDB: 1UC6) (*10*), LIF/gp130 D2D3 (PDB: 1PVH) (*11*), IL-6Rα D1-D3 (PDB: 1N26) (*12*), and IL6/IL-6Rα D2D3/gp130 D1-D3 (PDB: 1P9M) (*13*), as well as AlphaFold  predicted models of CLCF1, CRLF1, EBI3, p28, and IL27-Rα, were used as initial models for model building. A combination of selected initial models was docked into corresponding cryo-EM density map using Fit-in-map of UCSF Chimera (*4*). The models were adjusted manually in Coot (*1*), followed by real space refinement in Phenix (*2*). For the highly flexible CNTF, CLCF1, LIF and IL-27 complexes, the well resolved regions around the interaction core of each complex were manually modeled and refined. Models containing full ECDs of each signaling complex were also generated by combining the structures of the interaction core region and models of the receptor distal domains, which were derived from published structures (gp130 D1, PDB: 1P9M; gp130 D6, PDB: 3L5H; gp130 D4-D6, PDB: 3L5H; LIFR D1, PDB:3E0G), a structure obtained in this study (CNTFRα D1, Figure S2), or AlphaFold predicted models (LIFR D6-D8 and IL-27Rα D3-D5). The composite models with full ECDs were refined against each full cryo-EM map using rigid body refinement in Phenix. The manually built and refined structures of the complex interaction core region were further subjected to PDBePISA analysis (*14*) to calculate buried surface areas of complex interfaces. Full models containing the entire ECDs were used to show the overall architectures of the complexes and analyze relative positions of the membrane-proximal domains. All structural figures were made in UCSF Chimera.

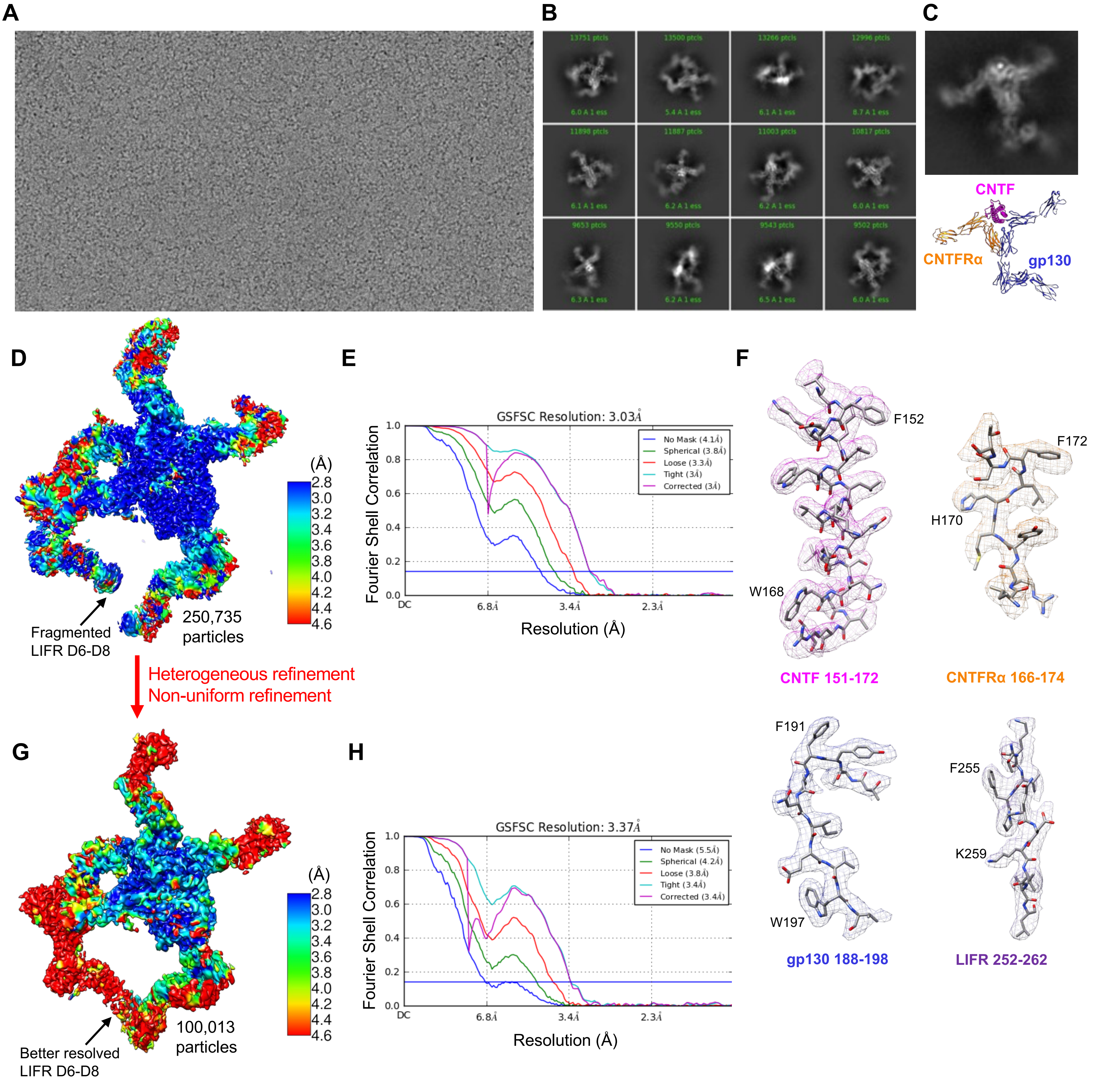


**Fig. S1. Cryo-EM analysis of the CNTF signaling complex**

(A-B) Representative raw micrograph and 2D class averages.
(C) CNTF complex intermediate (CNTF/CNTFRα/gp130 sub-complex) observed on the EM grid.
(D) Local resolution estimation of the final cryo-EM density map showing around 2.8 Å local resolution at the interaction core region. This map was used for building the atomic model of the interaction core region including CNTF, CNTFRα D2D3, gp130 D2-D5, and LIFR D2-D5. However, the map has fragmented LIFR D6-D8 density due to flexibility-induced heterogeneity of this region.
(E) FSC curve of the CNTF complex reconstruction showing a global resolution of 3.03 Å with the 0.143 gold standard threshold.
(F) Representative cryo-EM density of the 3D reconstruction in (D).
(G) The 250,735 particles from (D) were further subjected to heterogeneous refinement, identifying 100,013 particles with more homogeneous LIFR D6-D8. Non-uniform refinement of this subset of particles yielded a map with better density at LIFR FNIII domains. The local resolution of the map was estimated as in (D), showing lower global resolution. This map was used to generate a complete model of the CNTF complex with full ECDs by combining the structure of the interaction core region and models of the receptor distal domains derived from published structures (gp130 D1, PDB: 1P9M; gp130 D6, PDB: 3L5H; LIFR D1, PDB:3E0G), a structure obtained in this study (CNTFRα D1, fig. S2), and a model predicted by AlphaFold (LIFR D6-D8).
(H) FSC curve of the CNTF complex reconstruction in (G) showing a global resolution of 3.37 Å with the 0.143 gold standard threshold.

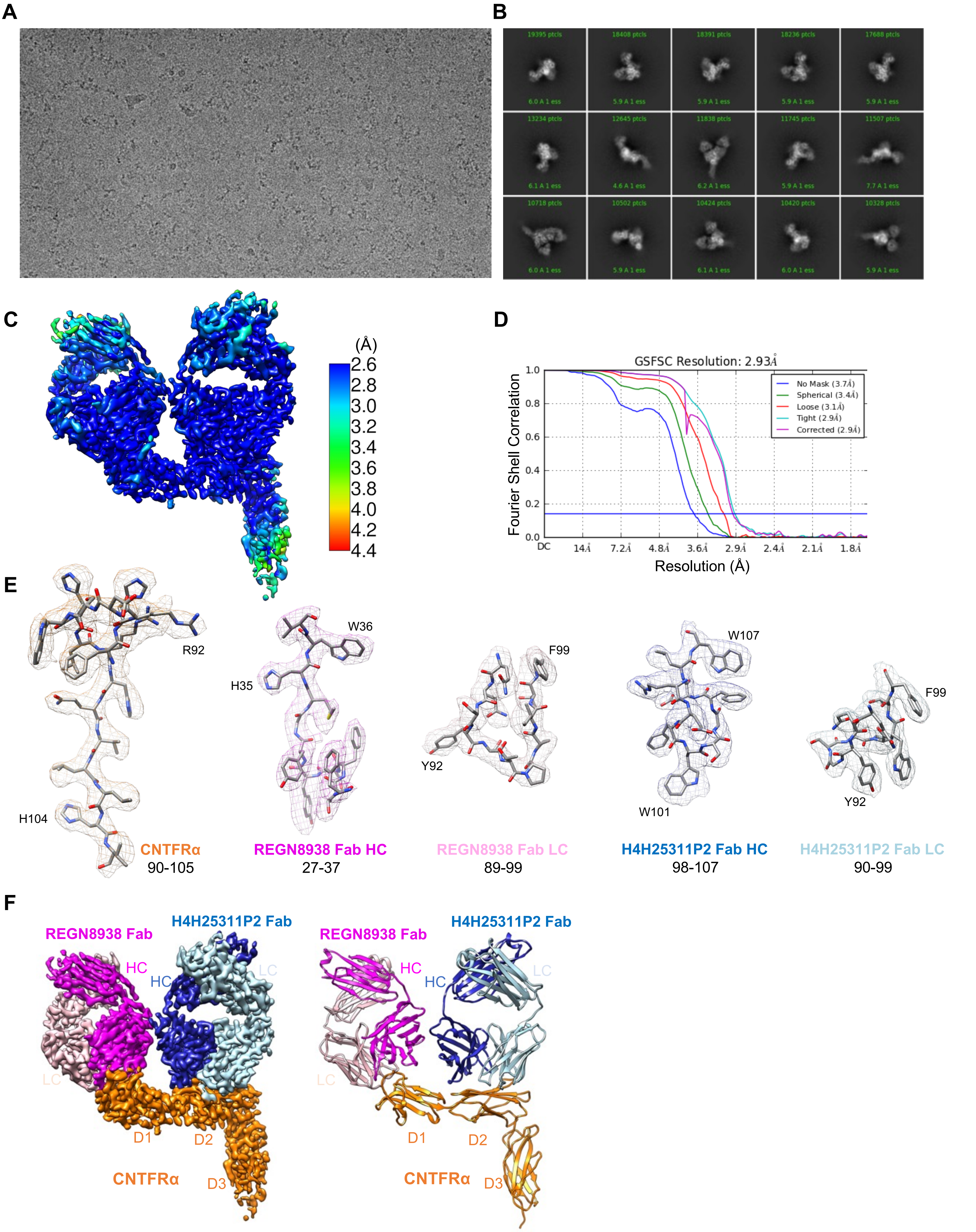


**Fig. S2. Cryo-EM structure of CNTFRα in complex with REGN8938 Fab and H4H25322P2 Fab**

(A-B) Representative raw micrograph and 2D class averages.
(C) Local resolution estimation of the final cryo-EM density map showing better than 3 Å resolution for the majority of CNTFRα.
(D) FSC curve of the CNTFRα/REGN8938 Fab/H4H25322P2 Fab complex reconstruction showing a global resolution of 2.93 Å with the 0.143 gold standard threshold.
(E) Representative cryo-EM density of the complex. HC: heavy chain; LC: light chain.
(F) Colored density map and atomic model of the CNTFRα/REGN8938 Fab/H4H25322P2 Fab complex. HC: heavy chain; LC: light chain.

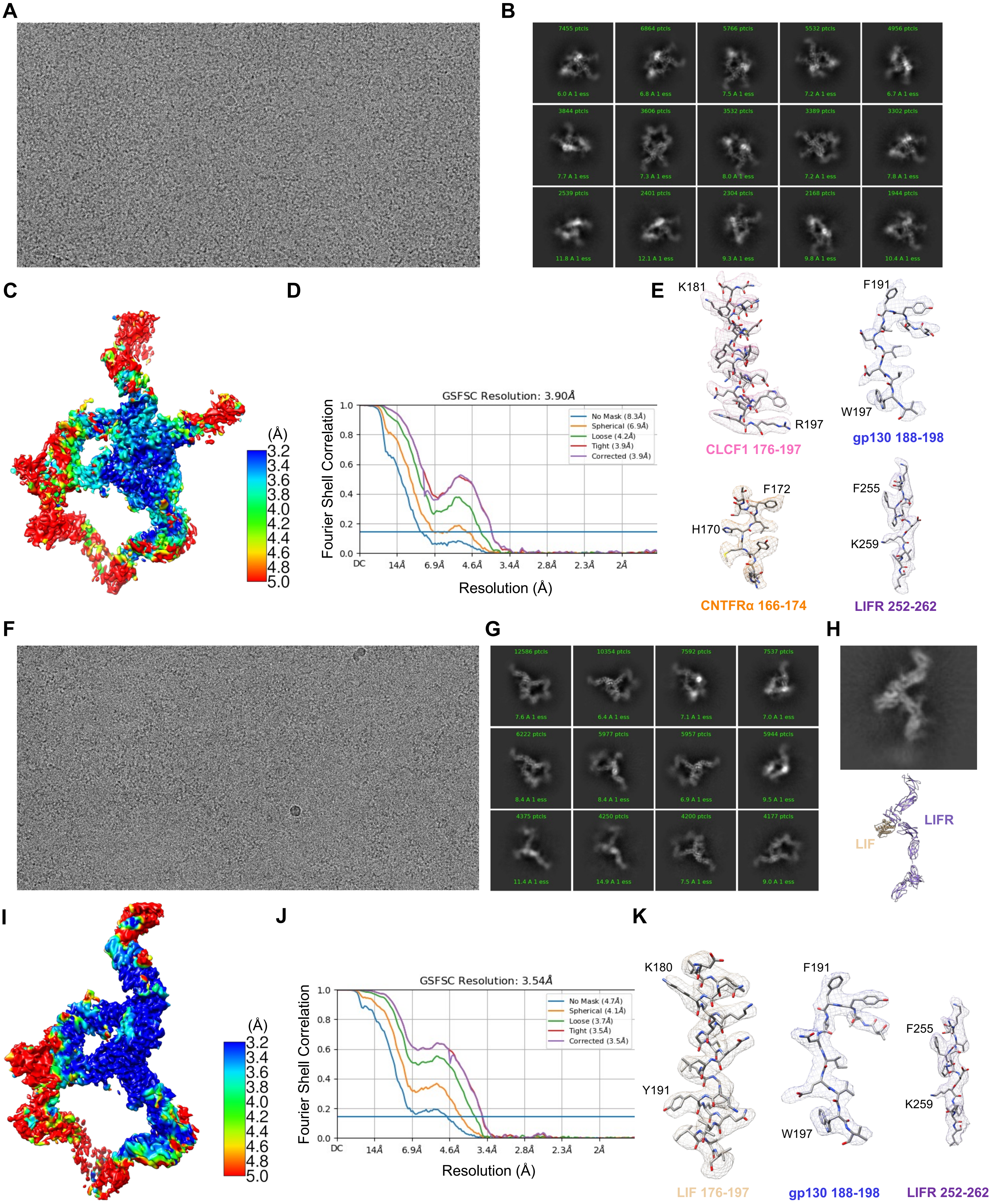

A
B
C
D
E
F
G
H
I
J
K
GSFSC Resolution: 3.90Å
Fourier Shell Correlation
Resolution (Å)
No Mask (8.3Å)
Spherical (6.9Å)
Loose (4.2Å)
Tight (3.9Å)
Corrected (3.9Å)
K181
R197
CLCF1 176-197
F191
W197
gp130 188-198
F172
H170
CNTFRα 166-174
F255
K259
LIFR 252-262
LIFR
LIF
GSFSC Resolution: 3.54Å
No Mask (4.7Å)
Spherical (4.1Å)
Loose (3.7Å)
Tight (3.5Å)
Corrected (3.5Å)
K180
Y191
LIF 176-197
gp130 188-198
LIFR 252-262


Figure S3

**Fig. S3. Cryo-EM analysis of the CLCF1 signaling complex and the LIF signaling complex**

(A-B) Representative raw micrograph and 2D class averages of the CLCF1 signaling complex .
(C) Local resolution estimation of the final cryo-EM density map of the CLCF1 complex showing around 3.4 Å local resolution at the interaction core region, including CLCF1, CNTFRα D2D3, gp130 D2-D5, and LIFR D2-D5.
(D) FSC curve of the CLCF1 complex reconstruction showing a global resolution of 3.90 Å with the 0.143 gold standard threshold.
(E) Representative cryo-EM density of the 3D reconstruction in (C).
(F-G) Representative raw micrograph and 2D class averages of the LIF signaling complex .
(H) LIF complex intermediate (LIF/LIFR sub-complex) observed on the EM grid.
(I) Local resolution estimation of the final cryo-EM density map showing around 3.2 Å local resolution at the interaction core region, including LIF, gp130 D2-D5, and LIFR D2-D5.
(J) FSC curve of the LIF complex reconstruction showing a global resolution of 3.54 Å with the 0.143 gold standard threshold.
(K) Representative cryo-EM density of the 3D reconstruction in (I).

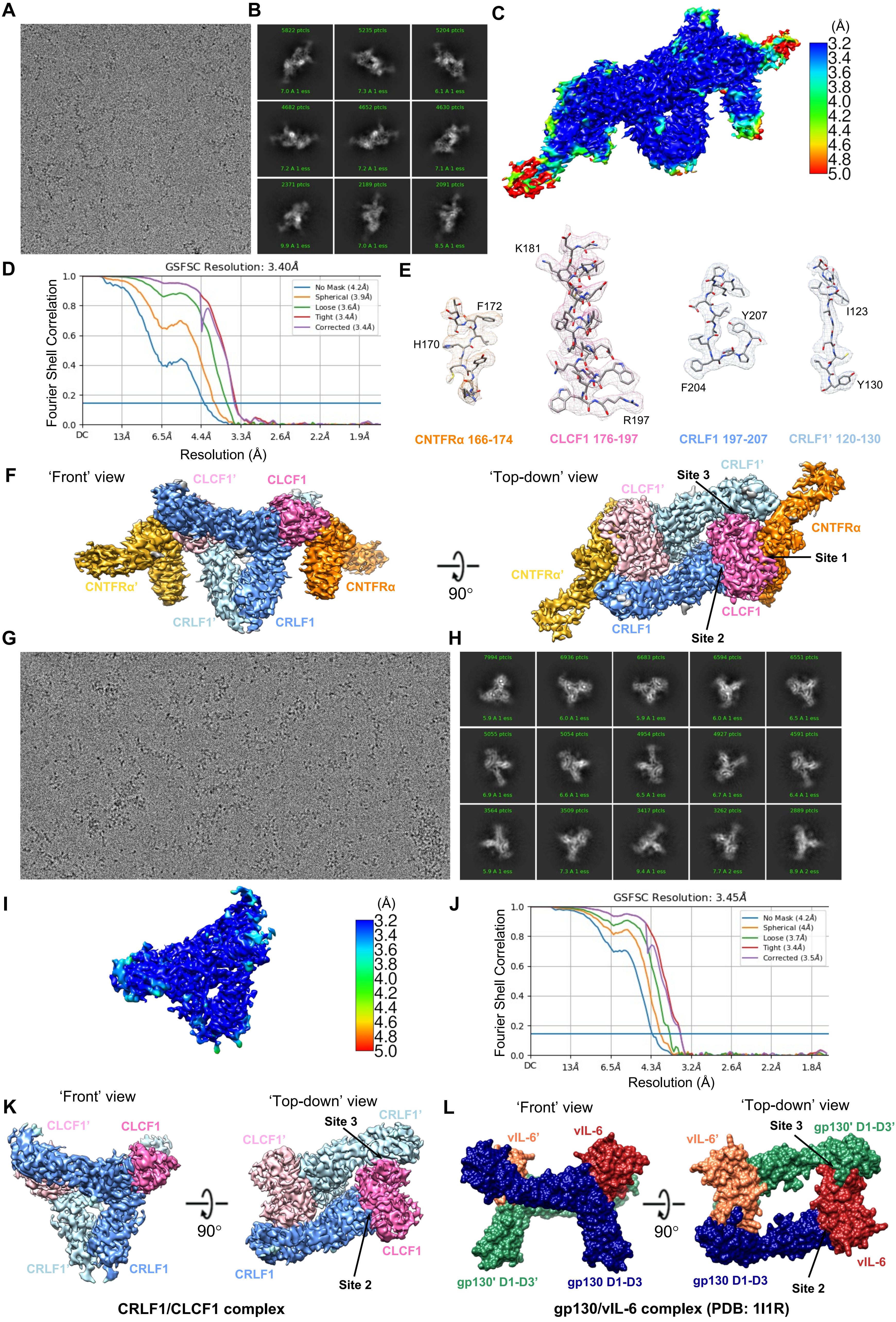
A
B
C
(Å)
3.2
3.4
3.6
3.8
4.0
4.2
4.4
4.6
4.8
5.0
D
GSFSC Resolution: 3.40Å
Fourier Shell Correlation
No Mask (4.2Å)
Spherical (3.9Å)
Loose (3.6Å)
Tight (3.4Å)
Corrected (3.4Å)
DC
13Å
6.5Å
4.4Å
3.3Å
2.6Å
2.2Å
1.9Å
Resolution (Å)
E
K181
F172
H170
Y207
I123
F204
Y130
R197
CNTFRα 166-174
CLCF1 176-197
CRLF1 197-207
CRLF1' 120-130
F
'Front' view
CLCF1'
CLCF1
CNTFRα'
CNTFRα
CRLF1'
CRLF1
90°
'Top-down' view
Site 3
Site 1
Site 2
G
H
I
(Å)
3.2
3.4
3.6
3.8
4.0
4.2
4.4
4.6
4.8
5.0
J
GSFSC Resolution: 3.45Å
Fourier Shell Correlation
No Mask (4.2Å)
Spherical (4Å)
Loose (3.7Å)
Tight (3.4Å)
Corrected (3.5Å)
DC
13Å
6.5Å
4.3Å
3.2Å
2.6Å
2.2Å
1.8Å
Resolution (Å)
K
'Front' view
'Top-down' view
CLCF1'
CLCF1
CRLF1'
CRLF1
Site 3
Site 2
90°
CRLF1/CLCF1 complex
L
'Front' view
'Top-down' view
vIL-6'
vIL-6
gp130' D1-D3'
gp130 D1-D3
Site 3
Site 2
90°
gp130/vIL-6 complex (PDB: 1I1R)

Figure S4

**Fig. S4. Cryo-EM analysis of the CRLF1-CLCF1-CNTFRα complex and the CRLF1-CLCF1 complex**

(A-B) Representative raw micrograph and 2D class averages of the CRLF1-CLCF1-CNTFRα complex.
(C) Local resolution estimation of the final cryo-EM density map of the CRLF1-CLCF1-CNTFRα complex.
(D) FSC curve of the CRLF1-CLCF1-CNTFRα complex reconstruction showing a global resolution of 3.40 Å with the 0.143 gold standard threshold.
(E) Representative cryo-EM density of the 3D reconstruction in (C).
(F) Colored density map of the CRLF1-CLCF1-CNTFRα complex in 'front' and 'top-town' views showing 2-fold symmetry of the complex. The two sets of molecules in the hexameric complex are annotated as CRLF1, CLCF1, CNTFRα, CRLF1', CLCF1', and CNTFRα'.
(G-H) Representative raw micrograph and 2D class averages of the CRLF1-CLCF1 complex .
(I) Local resolution estimation of the final cryo-EM density map of the CRLF1-CLCF1 complex .
(J) FSC curve of the CRLF1-CLCF1 complex reconstruction showing a global resolution of 3.45 Å with the 0.143 gold standard threshold.
(K) Colored density map of the CRLF1-CLCF1 complex in 'front' and 'top-town' views showing 2-fold symmetry of the complex. The two sets of molecules in the tetrameric complex are annotated as CRLF1, CLCF1, CRLF1', and CLCF1'.
(L) Surface representation of the gp130 D1-D3/vIL-6 complex crystal structure (PDB: 1I1R) in 'front' and 'top-town' views showing similar architectures of this complex and the CRLF1-CLCF1 complex.

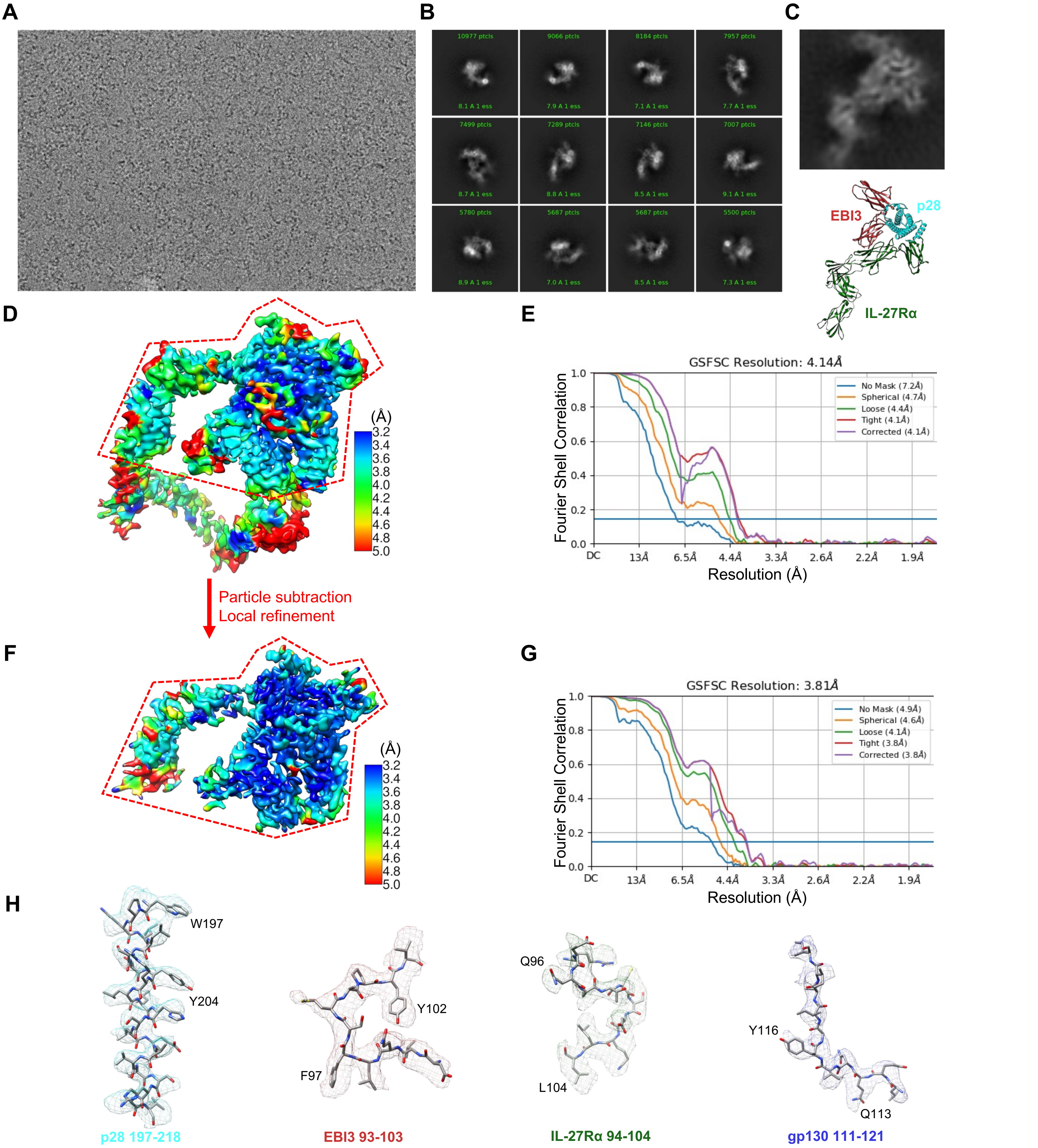


**Fig. S5. Cryo-EM analysis of the IL-27 signaling complex**

(A-B) Representative raw micrograph and 2D class averages.
(C) IL-27 complex intermediate (p28/EBI3/IL-27Rα sub-complex) observed on the EM grid.
(D) Local resolution estimation of the final cryo-EM density map showing around 3.6 Å local resolution at the interaction core region. This map was used to build a complete model of the IL-27 signaling complex. Models of gp130 D4-D6 derived from PDB:3L5H and IL-27Rα D3-D5 predicted by AlphaFold were rigid-body refined against the EM density.
(E) FSC curve of the full IL-27 complex reconstruction showing a global resolution of 4.14 Å with the 0.143 gold standard threshold.
(F) Particles from (D) was subjected to particle subtraction and focused refinement around the interaction core region, including p28, EBI3, IL-27Rα D1D2, and gp130 D1-D3. Local resolution estimation of the focused refinement map shows around 3.4 Å local resolution at the binding interfaces. This map was used to build the atomic model for the interaction core region including p28, EBI3, gp130 D1-D3, and IL-27Rα D1D2.
(G) FSC curve of the focused refinement map showing a resolution of 3.81 Å with the 0.143 gold standard threshold.
(H) Representative cryo-EM density of the 3D reconstruction in (F).

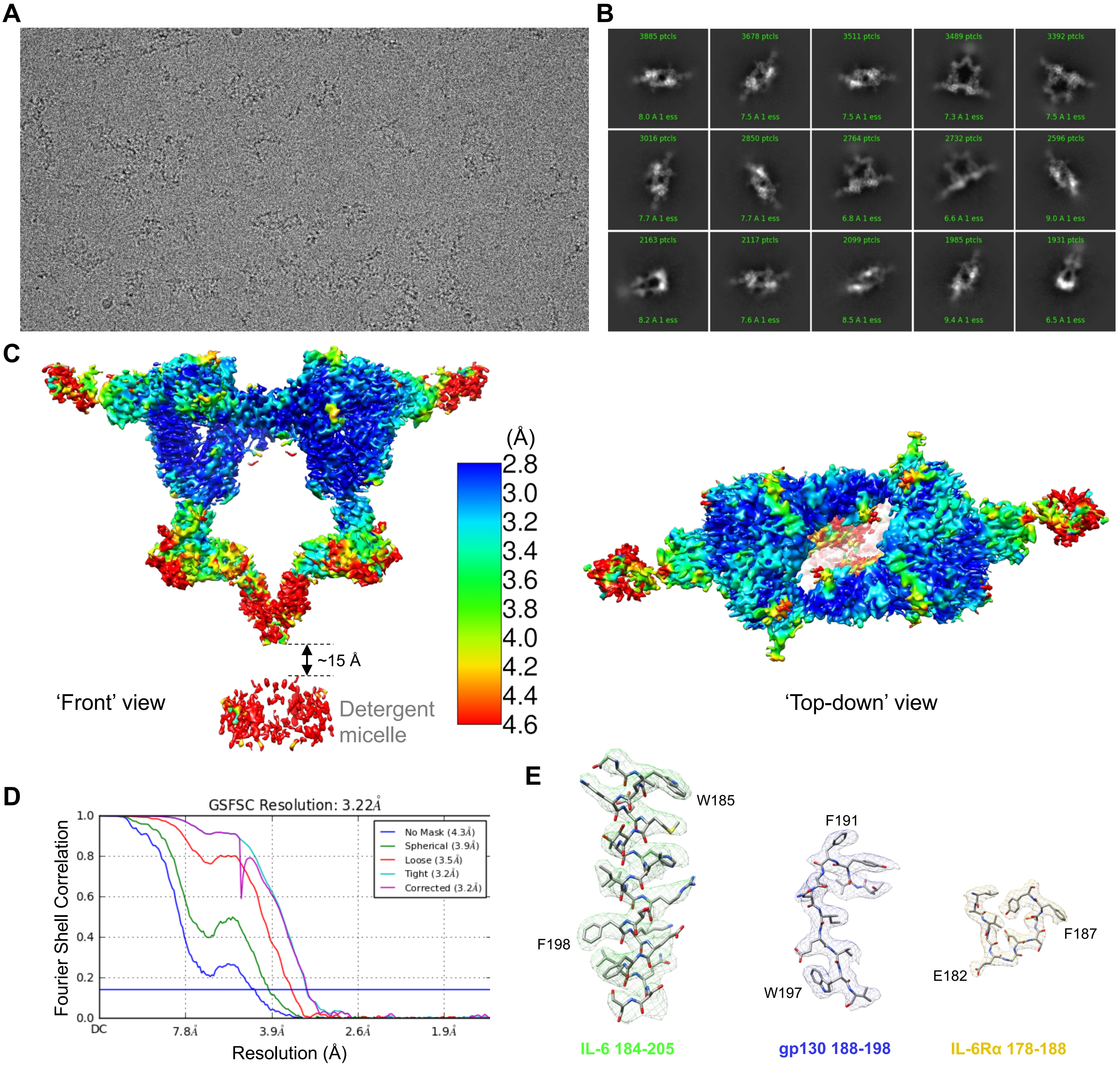


**Fig. S6. Cryo-EM analysis of the IL-6 signaling complex in detergent**

(A-B) Representative raw micrograph and 2D class averages.
(C) Local resolution estimation of the final cryo-EM density map in 'front' and 'top-town' views showing around 2.8 Å local resolution at the interaction core region and 2-fold symmetry of the complex. The TM domain of gp130 is not resolved in the detergent micelle. There is a ~15 Å gap between gp130 C-terminal density and the detergent micelle.
(D) FSC curve of the IL-6 complex reconstruction showing a global resolution of 3.22 Å with the 0.143 gold standard threshold.
(E) Representative cryo-EM density of the 3D reconstruction in (C).

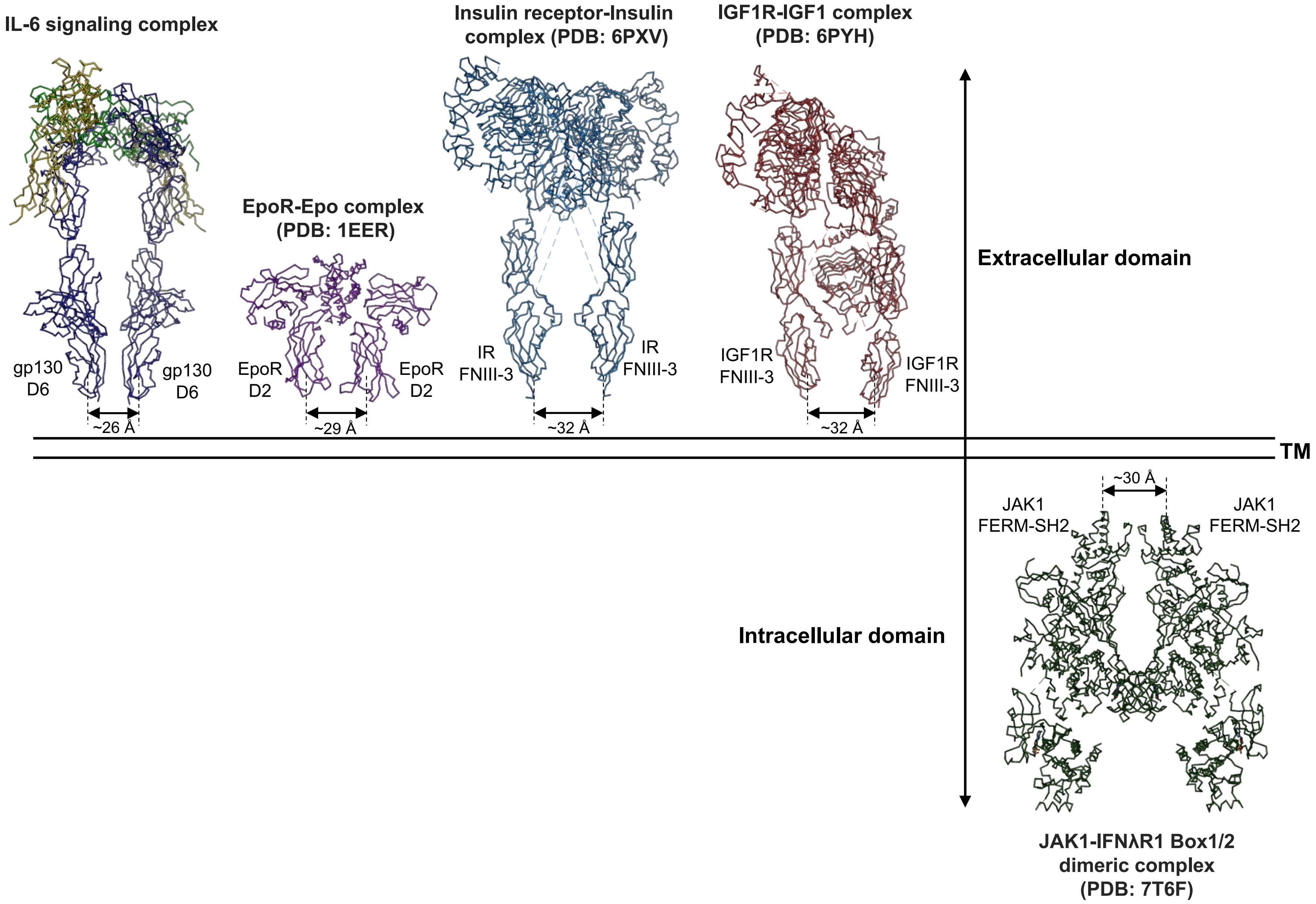


**Fig. S7. Comparison of distances between membrane-proximal domains in various cytokine signaling complexes and the intracellular JAK1-IFNλR1 dimeric complex**

The distances between the bottom centers of the membrane-proximal domains of the two signaling receptors in the IL-6 complex, EpoR-Epo complex (PDB: 1EER), Insulin receptor-insulin complex (PDB: 6PXV), IGF1R-IGF1 complex (PDB: 6PYH) are all around 30 Å. Moreover, on the intracellular side, the distance between the two membrane-proximal FERM-SH2 domains in the JAK1-IFNλR1 dimeric complex (PDB: 7T6F) is also about 30 Å.

**Table S1. Sequence identity scores (%) among six gp130 family cytokines**

| | CNTF | CLCF1 | LIF | IL-27 p28 | IL-6 | OSM |
|---|---|---|---|---|---|---|
| **CNTF** | 100 | 13.9 | 10.9 | 13 | 8.8 | 8.9 |
| **CLCF1** | 13.9 | 100 | 11.9 | 21.2 | 15.3 | 14.3 |
| **LIF** | 10.9 | 11.9 | 100 | 10 | 12.6 | 10.6 |
| **IL-27 p28** | 13 | 21.2 | 10 | 100 | 14.8 | 12.5 |
| **IL-6** | 8.8 | 15.3 | 12.6 | 14.8 | 100 | 8.6 |
| **OSM** | 8.9 | 14.3 | 10.6 | 12.5 | 8.6 | 100 |

**Table S2. Cryo-EM data collection, processing, and refinement statistics, related to STAR Methods**

| | CNTF signaling complex | | CLCF1 signaling complex | | LIF signaling complex | | IL-27 signaling complex | | IL-6 signaling complex | CRLF1-CLCF1-CNTFRα complex | CNTFRα-REGN8938 Fab-H4H25322P2 Fab complex |
|---|---|---|---|---|---|---|---|---|---|---|---|
| **Data collection and processing** | | | | | | | | | | | |
| Magnification | 105,000 | | 105,000 | | 105,000 | | 105,000 | | 105,000 | 105,000 | 105,000 |
| Voltage (kV) | 300 | | 300 | | 300 | | 300 | | 300 | 300 | 300 |
| Electron exposure ($e^-$/Å$^2$) | 40 | | 40 | | 40 | | 40 | | 40 | 40 | 40 |
| Defocus range (μm) | -1.4 to -2.6 | | -1.4 to -2.6 | | -1.4 to -2.6 | | -1.4 to -2.6 | | -1.4 to -2.6 | -1.4 to -2.6 | -1.4 to -2.6 |
| Pixel size (Å) | 0.85 | | 0.86 | | 0.86 | | 0.86 | | 0.85 | 0.86 | 0.85 |
| Number of movies | 6,511 | | 25,442 | | 11,007 | | 26,610 | | 12,143 | 9,943 | 7,859 |
| Initial number of particles | 3,197,974 | | 13,119,612 | | 5,545,456 | | 19,480,836 | | 3,881,621 | 1,258,062 | 4,515,671 |
| Final selected particles | 250,735 | 100,013 | 92,463 | | 171,328 | | 139,752 | | 105,760 | 117,773 | 568,328 |
| Symmetry imposed | C1 | | C1 | | C1 | | C1 | | C2 | C2 | C1 |
| Map resolution (Å) | 3.03 | 3.37 | 3.90 | | 3.54 | | 3.81 | 4.14 | 3.22 | 3.40 | 2.93 |
| FSC threshold | 0.143 | | 0.143 | | 0.143 | | 0.143 | | 0.143 | 0.143 | 0.143 |
| **Refinement** | | | | | | | | | | | |
| Map sharpening *B* factor (Å$^2$) | -70 | -49.6 | -90 | | -51.4 | | -120 | -90 | -78 | -94.5 | -99.6 |
| Model composition | | | | | | | | | | | |
| Non-hydrogen atoms | 9,640 | 14,639 | 9,628 | 14,642 | 7,912 | 12,267 | 7,101 | 11,419 | 17,366 | 12,542 | 8,960 |
| Protein residues | 1,163 | 1,793 | 1,167 | 1,799 | 963 | 1,511 | 865 | 1,435 | 2,108 | 1,526 | 1,143 |
| Ligands | 18 | 18 | 16 | 16 | 15 | 15 | 11 | 11 | 38 | 32 | 8 |
| R.m.s. deviations | | | | | | | | | | | |
| Bond lengths (Å) | 0.002 | 0.002 | 0.002 | 0.002 | 0.002 | 0.002 | 0.002 | 0.002 | 0.002 | 0.002 | 0.002 |
| Bond angles (°) | 0.448 | 0.449 | 0.486 | 0.486 | 0.507 | 0.459 | 0.471 | 0.521 | 0.454 | 0.449 | 0.447 |
| Validation | | | | | | | | | | | |
| MolProbity score | 1.66 | 1.67 | 1.79 | 1.76 | 1.85 | 1.75 | 1.55 | 1.77 | 1.43 | 1.34 | 1.53 |
| Rotameric outliers (%) | 0.85 | 0.86 | 0.00 | 0.00 | 0.00 | 0.07 | 0.00 | 0.00 | 0.95 | 1.06 | 1.30 |
| Clash score | 5.34 | 5.94 | 5.56 | 5.84 | 6.62 | 6.45 | 4.91 | 6.75 | 3.18 | 2.71 | 5.45 |
| Ramachandran plot | | | | | | | | | | | |
| Favored (%) | 94.54 | 94.99 | 91.88 | 93.05 | 91.95 | 94.20 | 95.79 | 94.17 | 95.37 | 96.10 | 97.17 |
| Allowed (%) | 5.46 | 5.01 | 8.12 | 6.95 | 8.05 | 5.80 | 4.09 | 5.76 | 4.63 | 3.90 | 2.83 |
| Disallowed (%) | 0.00 | 0.00 | 0.00 | 0.00 | 0.00 | 0.00 | 0.12 | 0.07 | 0.00 | 0.00 | 0.00 |
| **Deposition ID** | | | | | | | | | | | |
| PDB | 8D74[*] | 8D7G[#] | 8D7R[*] | 8D7T[#] | 8D6A[*] | 8D70[#] | 8D85[*] | 8D83[#] | 8D82[#] | 8D7H | 8D7E |
| EMDB | 27227[a] | 27229[b] | 27231 | | 27221 | | 27247[c] | 27246[d] | 27244[e] | 27230 | 27228 |

[*] Structure of the complex interaction core region

[#] Structure of full extracellular domains (ECDs) of the complex

[a] Cryo-EM map of the CNTF signaling complex: refined from the selected full particle dataset, with better resolution in the interaction core region

[b] Cryo-EM map of the CNTF signaling complex: refined from a particle subset, with lower global resolution but improved LIFR D6-D8 density

[c] Cryo-EM map of the IL-27 signaling complex: focused refinement on the interaction core region

[d] Cryo-EM map of the IL-27 signaling complex with full ECDs

[e] Cryo-EM map of detergent-solubilized IL-6 signaling complex with full ECDs
